# A Rapid, High Throughput, Viral Infectivity Assay using Automated Brightfield Microscopy with Machine Learning

**DOI:** 10.1101/2022.03.23.485512

**Authors:** Rupert Dodkins, John R. Delaney, Tess Overton, Frank Scholle, Alba Frias, Elisa Crisci, Nafisa Huq, Ingo Jordan, Jason T. Kimata, Ilya G. Goldberg

## Abstract

Infectivity assays are essential for the development of viral vaccines, antiviral therapies and the manufacture of biologicals. Traditionally, these assays take 2–7 days and require several manual processing steps after infection. We describe an automated assay (AVIA™), using machine learning (ML) and high-throughput brightfield microscopy on 96 well plates that can quantify infection phenotypes within hours, before they are manually visible, and without sample preparation. ML models were trained on HIV, influenza A virus, coronavirus 229E, vaccinia viruses, poliovirus, and adenoviruses, which together span the four major categories of virus (DNA, RNA, enveloped, and non-enveloped). A sigmoidal function, fit to virus dilution curves, yielded an R^2^ higher than 0.98 and a linear dynamic range comparable to or better than conventional plaque or TCID_50_ assays. Because this technology is based on sensitizing AIs to specific phenotypes of infection, it may have potential as a rapid, broad-spectrum tool for virus identification.

---

Vaccination is the most efficacious and economic intervention for prevention of infectious diseases, spillover events from animals to humans, and (in the case of veterinary vaccines) food security ^1–3^. Importance of development of novel vaccines for pandemic preparedness, or adequate stockpiles of proven vaccines, has been drastically demonstrated in the coronavirus disease 2019 (COVID-19) outbreak that resulted in hundreds of millions of cases, and almost 6 million deaths worldwide to date^4^.

Research on vaccines and antiviral therapies depend on infectivity assays for quantification of vector design decisions, purification processes and efficacy. However, infectivity assays are also central to the development and release of biologicals, medicines made from living cells taken from animals and thus at risk of contamination with adventitious agents ^5,6^. As much as a third of all infectivity assays currently being done are for monitoring viral clearance in the manufacturing process for biological therapies ^7,8^. For such assays a set of standard viral stocks is used to spike a manufacturing process and the infectious titer is measured before and after various process steps. The FDA outlines a reduction in infectivity of 10^15^ for all manufacturing steps combined, for 11 different viruses, and these assays are typically done with conventional plaque assays^9^.

All combined, any improvement in the pace of measuring viral infectivity, its cost, or the reliability of the result can thus have profound consequences for the development of conventional, lipid-encapsulated or viral-vectored vaccines, antiviral drugs and biocides, biological therapies that require clearance assays, as well as therapies that use viral vectors such as gene therapy, cancer immunotherapy, and oncolytic therapies.

Infectivity assays operate by observing and quantifying the effects of viral replication on a culture of susceptible or permissive cells. For example, in plaque assays, cells are infected on 6-well or 24-well plates and overlaid with an immobilizing medium that localizes viral spread to neighboring cells only ^10,11^. This causes infected cells to form localized clusters of cytopathic effects (CPEs; dead or fused cells) that can be identified and counted to measure viral infectivity. The problem is that it takes 2-14 days to form plaques large enough to be identified by eye ^12,13,14^. The long incubation time slows down research, the overlays interfere with automation, which limits throughput, and the large format of the assay contributes to costs, especially in high containment facilities^15^.

The Tissue Culture Infectious Dose (TCID_50_) assay is an endpoint dilution assay used to find a specific dilution of virus stock where 50% of the infected wells show signs of being infected after a set incubation period^16^. As with plaque assays, an extended incubation time is required for multiple rounds of infection to occur so that a well of cells initially infected with a single infectious viral particle will become visibly infected. Unlike plaque assays, the count of infection events is much lower, which contributes to reduced precision. The TCID_50_ assay is typically done on 96-well or 384-well plates, but the number of samples per plate is limited by the requirement for multiple wells (usually 6-8 per replicate) for each dilution^17^. For obvious infection phenotypes like cell lysis, a well can be identified as infected vs. uninfected reliably without additional reagents, while more subtle phenotypes or more reliable automation may require more expensive reagents to detect infection reliably. This assay can be largely automated and it provides an estimate of the titer range but it is not as precise as a standard plaque assay^18^. The limited sensitivity to small differences in infectivity can cause promising lead compounds or vaccines to be missed. Typically TCID_50_ assays determine titer to within a factor of 5 or 10, while plaque assays can be accurate to within a factor of 2 ^19,20^.

The fluorescent focus assay or focus forming assay (FFA) is a variation of the plaque assay, but instead of relying on multiple rounds of infection and cell lysis in order to detect plaque formation, FFA employs immunostaining with detection of particular viral antigen expressed in the infected cells and is developed either using fluorescently labeled antibodies or immunocytochemistry techniques^21^. Fluorescent protein genes can also be expressed by introducing them into a recombinant viral genome. FFA can be used to detect individual infected cells before a plaque is formed, allowing this technique to yield results in less time than plaque or TCID_50_ assays. The labeled infected cells can also be counted by flow cytometry ^22^. However, FFA is dependent on a consistent supply of antibody reagents or transgenic viral strains, and the experimental analysis requires multiple manipulations that can make the final quantification subjective. The requirement for reagents and processing steps necessary for antibody-based detection can be avoided by developing viral strains carrying genes for fluorescent proteins, luciferase or beta-galactosidase. Including marker genes in therapeutic viral vectors may be problematic, and not all viruses are available in recombinant formats^13^. Laser Force Cytology uses a flow cell and backscatter imaging to discern infected from uninfected cells ^23,24^. This technique possesses many of the benefits of FFA, such as rapid turnaround time, without the requirement of cell labeling. However, this method requires specialist hardware, has a low limit of detection, and has challenges in scalability across many samples due to its reliance on a flow cell. Using deep learning and convolutional neural networks has been previously used on brightfield images of infected cells to detect CPEs^27,28^. These studies focused on a single virus each rather than proposing a generalized infectivity platform, and unlike this study, the CPEs are manually visible at the timepoints that were used.

A key difference between AI-based image processing and conventional image analysis that is used in FFA processing or automated plaque counting, is how variability in the assay is accounted for. This includes variation in cell health and density, variation in virally-induced infection phenotypes, and variation from well to well, plate to plate, and day to day. Conventional image processing relies on choosing parameters that are sufficiently tolerant of the variability without adversely affecting the assay’s sensitivity or precision. The parameter choice is typically in the hands of the experimentalist, and this can make the assay results dependent on the parameters chosen or the person choosing them. This can be even more subjective when the assay is directly read out by the experimentalist without software. When using AI-based techniques, the variability present in the assay is trained into the AI, since there are no image processing parameters for the experimentalist to choose. While different experimentalists may introduce bias by consistently growing slightly different looking cells and producing slightly different viral infectivity phenotypes, this difference can be quantified by having the two researchers titer the same viral sample, and importantly, the AI can be trained to ignore these variations by being trained with samples from multiple researchers.

We present a novel infectivity assay that solves the shortcomings of the traditional approaches listed above. The automated viral infectivity assay (AVIA) uses computer vision methods based on machine learning (ML) to analyze brightfield images collected from 96-well plates infected with limited dilutions of virus-containing samples. AIs can be trained to detect subtle cell morphological differences associated with virus infection prior to any manually visible cytopathic effects like swelling, fusion, or cell death ^25,26,23^. This is made possible by these algorithms having access to the full 12-16 bits of dynamic range in modern sensors (as opposed to the approximately 6-7 bits of the human retina). These models detect trends across a vast range of images simultaneously and are specialized to ignore the large amounts of irrelevant image information in complex images such as brightfield. The use of brightfield microscopy obviates cell fixation and staining, which allows the technique to be more easily automated as well as making it faster, safer, more reproducible, and more cost effective. Due to the quantitative assay readout for each well, fewer dilutions are required per sample than TCID_50_, freeing up valuable plate real estate, which increases sample throughput and further reduces cost per sample. Furthermore, the assay does not rely on plaque formation and is therefore not limited to plaque forming viruses (see Supplementary Fig. 2 with HIV for example). Lastly, the increased automation, reduction of sample preparation steps, and fully automated analysis ensures that results are consistent from day to day and from researcher to researcher. Table 1 summarizes the key metrics of the described infectivity assays, where the values for the traditional techniques are compiled from publications that made these comparisons ^29,13,30,19,31, 32,33^.

**Tab. 1.** A summary of three traditional techniques of conducting a viral infectivity assay as well as the proposed AVIA technique. By eliminating many of the human intensive steps of the traditional techniques, AVIA enables greater reproducibility, scalability, and lower costs.

| Technique | Reproducibility | Time | Labor | Cost | Automation/<br>Scalability |
| --- | --- | --- | --- | --- | --- |
| AVIA (ours) | excellent | hours | low | low | high |
| Plaque Assay | good | days | high | low | low |
| TCID<sub>50</sub> | medium | days | high | medium -<br>high | high |
| FFA: Antibodies<br>(Ab) or GFP | medium | day | medium<sup>GFP</sup> -<br>high<sup>Ab</sup> | medium<sup>GFP</sup><br>- high<sup>Ab</sup> | low<sup>Ab</sup> -<br>medium<sup>GFP</sup> |

This paper presents the development and validation of the AVIA assay platform. In the Automated Viral Infectivity Assay section, we describe the assay and the metrics of success used in the investigations. The Viral versatility and Example inactivation assay sections showcase the training performance of AVIA on a selection of viruses investigated so far as well as an example inactivation assay. In Supplementary Materials, we explore optimization of training parameters, and generalizability of models across experimental conditions.

## Automated viral infectivity assay

The challenge in training machine learning models on images with changes that cannot be observed by eye is setting up experimental conditions so that the images used for training have a well-defined ground truth. To accomplish this, we start with the assumption that a successful infectious viral particle will induce a detectable morphological change in the host cell. This way, a binary classifier model could be trained using cells infected with a high multiplicity of infection (MOI) to serve as the positive class, and mock-infected cells with no exposure to viral particles to serve as the negative class. The MOI of the positive class must be sufficiently high that all cells can be assumed to be infected synchronously as any uninfected cells will contaminate the positive class with false positives and place a limit on the accuracy of the trained model. For example, for an MOI of 3, 1-e^-3^ = 95% of cells will be infected according to Poisson statistics. Several identical wells of infected and uninfected cells are imaged at multiple locations to satisfy the large number of images needed to train an AI model based on deep-learning/convolutional neural networks^34^.

The workflow for training an AVIA model is as follows – a cell culture is first plated and infected, just as in a traditional assay. The cells are then left to incubate while a series of brightfield images are captured with a high-throughput microplate imager ^35–38^. If the microplate imager includes an incubator (temperature control, CO_2_), then this can happen automatically, otherwise the plate has to be manually transferred to the microplate reader at the selected timepoints. Then, the images and plate layout are transferred to ViQi Inc. servers, at which point the processing steps are complete for the user.

Currently, when first operating on a new virus, cell line, or imaging device, a new AI needs to be trained using the same cells, virus, plates and imaging instrument as will be used routinely for infectivity assays (Fig. 1A). Once a model is created, the user can conduct various infectivity assays such as inactivation assays, clearance assays or routine virus titers (Fig. 1B). For a routine assay, a 96-well plate can contain 3-4 replicates and 3-4 virus dilutions, allowing 8-12 independent samples per 96-well plate.

**Fig. 1.**
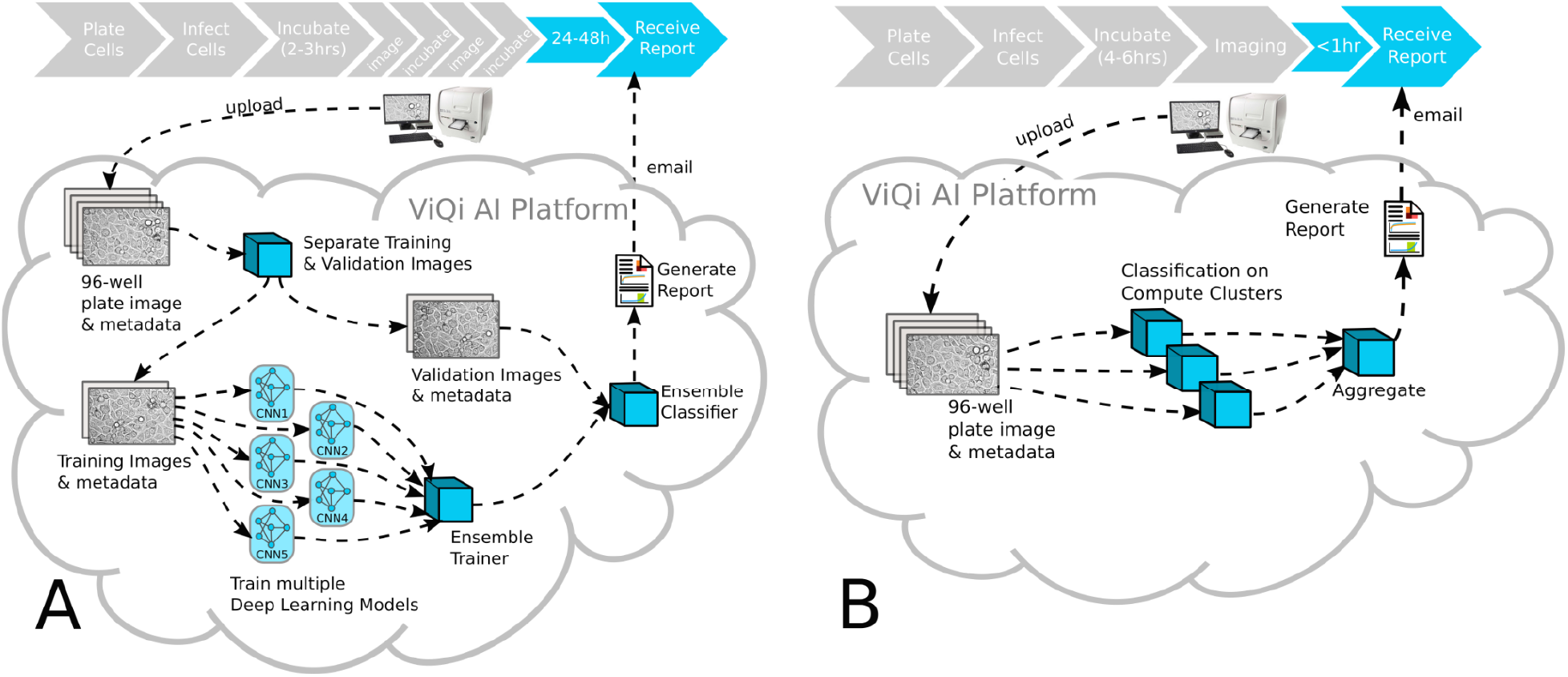
Model workflow for AVIA in training (A) and conducting an assay (B) using a trained model. After the cells are incubated, the remainder of the process is entirely automated improving reliability, cost, and scaling potential. The machine learning workflow includes training an ensemble of machine learning models to improve accuracy and generalizability.

Fig. 2 shows the input data for training and validating an AVIA model. A single 96-well plate is populated with a cell monolayer and infected at a range of MOIs according to the layout in Fig 2A, in which a quarter of the wells are infected at a high MOI and a quarter of the wells are uninfected (the training protocol), and half of the plate has a dilution series to validate the model performance on partially infected wells (validation protocol). To further increase the spatial resolution and maximize the number of training samples, the images are split up into square subregions, or ‘tiles’. Fig. 2B and 2C show example images from the two training quadrants and a potential tile separation scheme – here the images would each yield 256 tiles of size 128×128px.

**Fig. 2.**
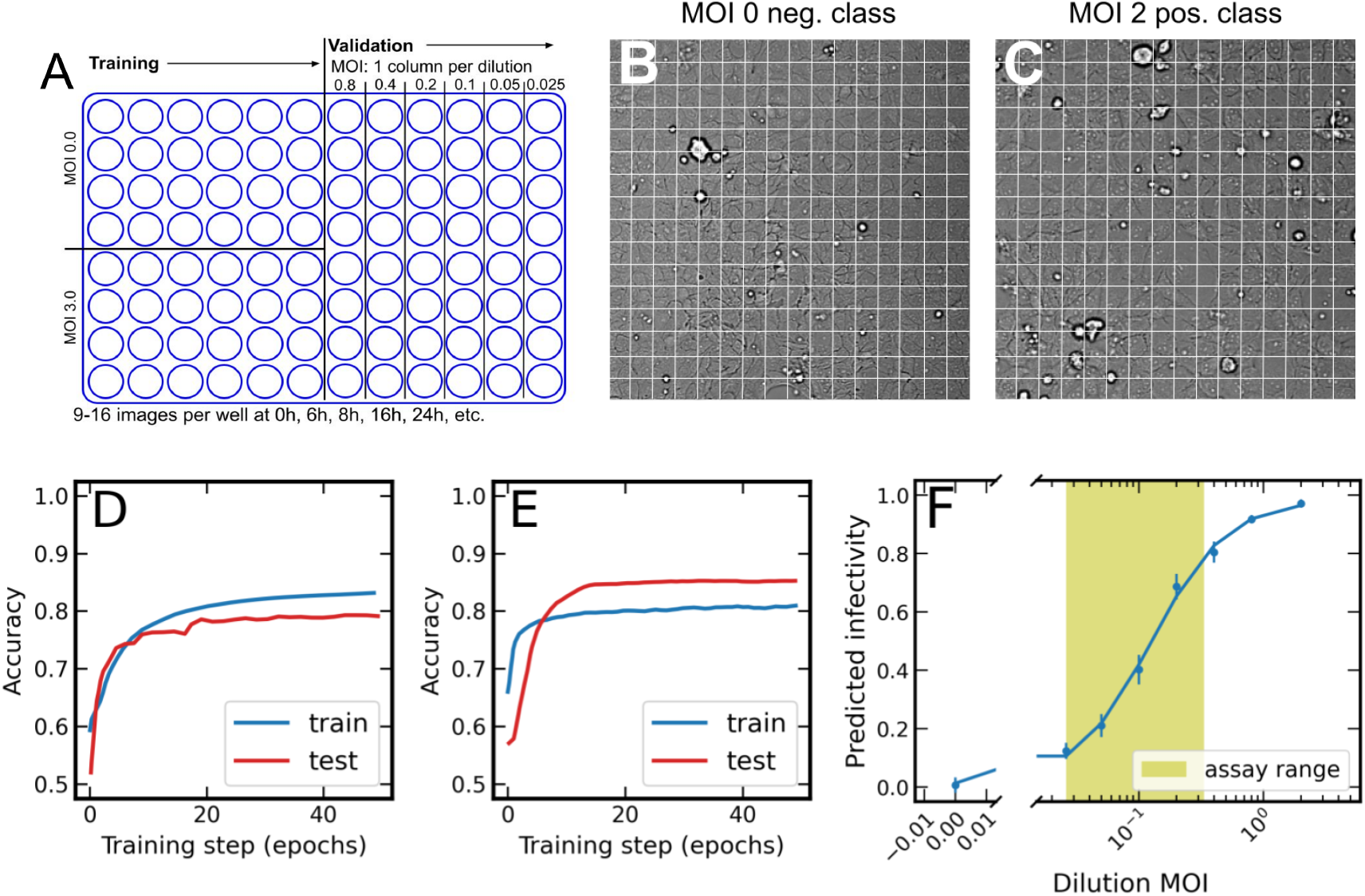
Training and validation of AVIA ML models. **A**, the plate is partitioned into training MOIs (mock infected and saturated infection) and validation MOIs (serial dilutions). **B** and **C**, brightfield images are split into tiles, which are used as inputs to train, test and validate the model. The performance of the model on test data relative to train tiles determines the degree of overtraining (**D** and **E**). **F**, a successful model predicts infectivity with plateaus at low and high MOI and a linear range at intermediate MOIs on validation data. The error bars show the standard error of the images for a given MOI.

While training, the model is exposed to many examples of images from either class, along with their labels, and the achieved binary accuracy is monitored for train (seen, blue) and test (unseen, red) data (Fig. 2D and E). When the test curve falls below the training curve, it demonstrates that the model is overtraining to the training data (Fig. 2D). The deep neural networks used in AVIA contain millions of free parameters, and with a finite amount of training data, some amount of overfitting to the specific examples in training is inevitable. Occasionally test data shows better performance than training data (Fig. 2E), indicating that the dropout and regularization training parameters used to suppress overtraining may be too aggressive.

A successfully generalized model would be able to identify tiles as infected or uninfected in images at a range of MOIs between the two extremes used in training. A sigmoidal trend is expected with plateaus towards the two extreme MOIs used in training and a linear range along which an infectivity assay would be sensitive (Fig. 2F). To quantify this, infectivity is defined as the fraction of tiles with a raw AI score above the threshold of 0.5 in a given image. These image scores are grouped according to MOI and their means and standard error define the predicted infectivity for a given MOI. This curve is fit with the four parameter logistic model (4PL)

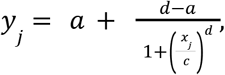

where y_j_ is the infectivity at MOI x_j_, *a* is the infectivity of the upper asymptote, *d* is the infectivity of the lower asymptote, *c* is the MOI at the inflection point of the curve, and b is related to the slope at the inflection point^39^. The datapoints from the training protocol (e.g. MOI 0.0 and 2.0), that were reserved for testing the AI, are included in these fits. To avoid using multiple points at the high and low plateaus in fitting Eq. 4PL, a series of fits were performed by cumulatively masking points from the high and low MOIs until the minimum valid number of datapoints was reached. The fits are evaluated by their R^2^ and the top 3 are reported. This curve also serves as the calibration for the assay, which allows converting the infectivity output of the AI (Y axis in Fig. 2F) to a multiplicity of infection (MOI) estimate. The quality of the fit indicates the quality of the predicted MOI when the AI’s infectivity output falls within the linear range of the assay (green zone in Figure 2F).

The kinetics of each virus and cell line combination vary. It is not clear *a priori* when the ideal time is to train and conduct an assay after infection. As time progresses, the signals from infection will generally get stronger, until they are so strong that they can be observed by eye. There is therefore an optimization of choosing a timepoint as early as possible to minimize the turnaround time of model training and assaying, but still retaining sufficient accuracy that the results are trustworthy and interpretable. Training the AI on phenotypes associated with productive infection rather than cell death or necrosis should also help with specificity. We therefore train models at several timepoints during the infection and compare the results. These experiments can consist of several binary models that then get compared or one multiclass model with infected and uninfected classes for the entire timecourse. This time analysis is typically performed before training and validating at a single timepoint.

## Results

### Viral versatility

With the model workflow optimized, and the degree of model generalizing power quantified, we are now in a position to demonstrate the application of AVIA on a set of viruses. For each virus, a separate model was trained using the MobileNet architecture, 128px tiles, 40% dropout, 1e-4.5 learning rate, but different selected assay timepoint and number of base CNNs that compromise the model – single (S), 8 CNNs (E8), or 16 CNNs (E16). These performances could be improved by gathering more training examples, exploring more of the hyperparameter space, or incorporating more hyperparameter combinations into an ensemble model architecture if necessary. The assay timepoint was determined by selecting the minimum timepoint in the productive phase of the infection (i.e., not the first timepoint) with the highest test accuracy. This timepoint was used for training binary AIs for each virus (infected vs. uninfected), and validating their predictions using serial dilutions with the indicated MOIs determined by conventional means (plaque assay or TCID_50_).

Fig. 3 shows the results of training and validation for each of the seven viruses. For each column, the top panel shows the development of the model performance on training and testing data as the training progresses through its cycles (epochs). The middle panel separates test performance into infected and uninfected classes to reveal any biases in the model towards false positives or false negatives. The bottom panel demonstrates model performance on MOIs from serial dilutions.

**Fig. 3.**
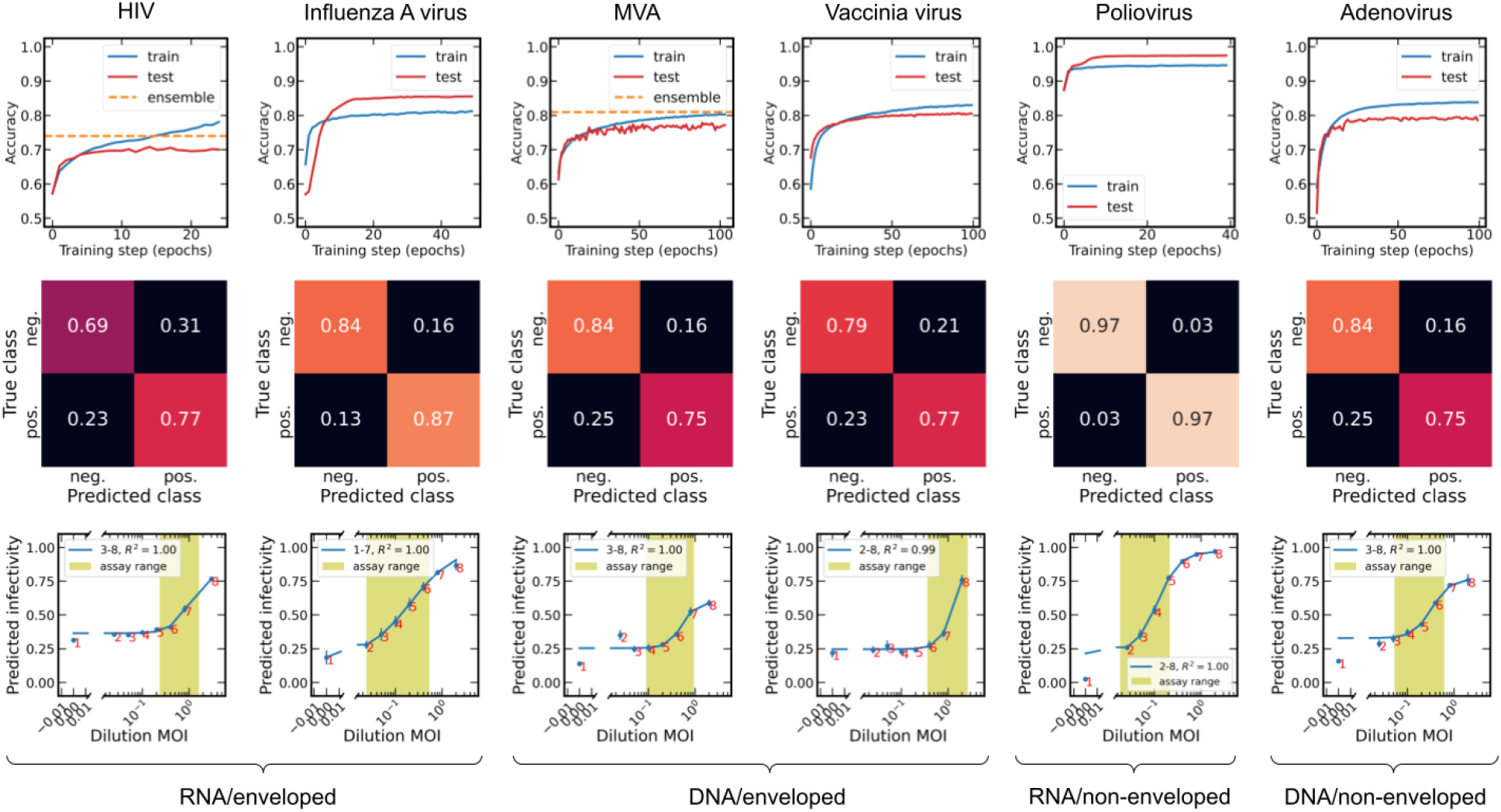
Model training and validation performance on six viruses. The top row shows the progress of model accuracy on train (seen by the AI, blue) and test (unseen by the AI, red) data, the middle row shows the test data accuracy for each of the two classes, the bottom row shows the model predictions on a range of intermediate MOIs from serial dilutions. The green box defines the assay’s linear range from the best fit for a sigmoidal function as described in the text. The error bars show the standard error of the images for a given MOI.

After training hundreds of models on different experimental setups and hyperparameter conditions a few metrics of success have been developed for model training and validation. These are:

1. A test accuracy plateau (or subsequent downtrend after peak accuracy) within the range of training epochs.
2. A peak test accuracy above 75%.
3. A peak test accuracy greater than 95% of the peak training accuracy.
4. A ratio of true positive rate to true negative rate (and vice versa) of less than 1.10.
5. A linear range of the assay, as determined by the sigmoid fit, within the bounds defined by the MOI dilutions in the calibration curve.
6. A linear range spanning more than a 10-fold range (comparable to that used in a traditional plaque assay with a countable range of 10-100 plaques per well) or equivalently containing ≥ 3 MOI dilutions.
7. A difference in the predicted MOI plateaus of ≥ 0.5.

Applying these seven metrics to the seven virus models shown, it is observed that all models satisfy criteria #1 and #2. These models were all trained with the EfficientNet-based CNN architecture which has smooth monotonic curves on AVIA data tested, as opposed to the VGG-based architecture that tended to find multiple local minima per 100 epochs for the selected hyperparameters. HIV would have failed criteria #3, but with the aid of the ensemble model method, all models pass. Three models narrowly fail criteria #4 achieving a ratio of 1.12 for HIV, MVA and adenovirus.

All but one model (vaccinia) satisfy criteria #5. This criteria is meant to ensure that the upper and lower bounds of the linear range are not beyond the MOIs in the calibration curve. Vaccinia appears to not have a well defined upper bound. This is either caused by an overestimate of the virus titer, or could potentially be addressed by improving the AI accuracy, thereby improving the false positive rate and shifting the calibration curve to the left. An improvement can be achieved using an ensemble AI or potentially larger tiles, which is currently under investigation.

Three models fail criteria #6, and have an assay range < 10-fold. In two cases (HIV and adenovirus), their linear ranges are still enough to cover three 2-fold dilutions for an 8-fold assay range. Vaccinia has a reduced range of 6.7. This can be addressed experimentally in the assays by using 2 or 3-fold dilutions rather than 5 or 10-fold ones. The AI can also be potentially improved as discussed above. Lastly, two models fail criteria #7. HIV has a linear prediction range of 0.5, passing narrowly, and the two MVA strains have a range of 0.4. Both MVA strains were already using a 16-model ensemble, so further improvements to AI accuracy could potentially come from larger tiles, or using both larger tiles and dilution data for training regressor AIs (see discussion). A reduced numerical range for valid predictions doesn’t appear to adversely affect the other aspects of assay performance discussed above, however.

Table 2 shows the versatility of AVIA. It demonstrates that a model could be trained for a virus/cell line combination within each major virus category of RNA, DNA, enveloped, and non-enveloped. Each model’s performance on dilution validation data achieves a clear sigmoidal trend correlated with MOI with an R^2^ > 0.99. These AI models were captured on four different microplate reader models, at five different laboratories, on seven different cell lines.

**Tab. 2.** Model performances for viruses in AVIA assays. Max AI accuracy is the test accuracy at the peak of the training curve. The linear assay range (MOI) is the difference in predicted MOI between where the linear range meets the upper lower asymptotes. Corona 229E has no corresponding validation data.

| Virus | Virus type | Assay | Max AI | Single model | Linear assay | Predicted | Sigmoid |
| --- | --- | --- | --- | --- | --- | --- | --- |

**Tab. 2 | Model performances for viruses in AVIA assays.**
|  |  | timepoint | accuracy | or Ensemble | range (MOI) | MOI range | R<sup>2</sup> |
| --- | --- | --- | --- | --- | --- | --- | --- |
| Coronavirus 229E | RNA/enveloped | 4h | 80% | S | N/A | N/A | N/A |
| HIV | RNA/enveloped | 24h | 74% | E8 | 0.2 - 1.6 | 0.4 - 0.9 | 0.996 |
| Influenza A virus | RNA/enveloped | 10h | 89% | S | 0.03 - 0.5 | 0.2 - 0.9 | 0.973 |
| MVA-CR | DNA/enveloped | 4h | 81% | E16 | 0.1 - 0.9 | 0.3 - 0.7 | 0.990 |
| MVA-WT | DNA/enveloped | 4h & 6h | 73% | E16 | 0.2 - 1.6 | 0.4 - 0.8 | 1.000 |
| Vaccinia | DNA/enveloped | 24h | 78% | S | 0.3 - 2.0 | 0.3 - 0.9 | 0.995 |
| Polio | RNA/<br>non-enveloped | 21h | 97% | S | 0.02 - 0.2 | 0.2 - 1.0 | 1.000 |
| Adenovirus | DNA/<br>non-enveloped | 24h | 87% | S | 0.01 - 0.3 | 0.3 - 0.8 | 0.996 |

### Example inactivation assay

The primary goal of AVIA is to create a viable method of infectivity quantification to increase the rate of vaccine development and other assays dependent on quantifying live virus titers. In this section, we demonstrate the applicability of AVIA in the context of an inactivation assay. Specifically, the model trained on Influenza A virus-infected MDCK cells at 16h after infection was used to quantify the effect of a biocide on infectivity of this virus.

The virus sample used in this experiment was treated with either media or the biocide product. The cells were then infected at a range of dilutions of virus, incubated, and imaged at 24h, 30h and 48h. If the biocide was successful, the dilution curve would be translated left towards lower MOIs.

Fig. 4 shows the AVIA predicted dilution curves for the treated and untreated populations at the three timepoints after infection. The same sigmoidal four-parameter model was used to fit treated and untreated dilutions at each timepoint. One of the four parameters, the inflection point, marks the location where the slope changes direction. The difference in the inflection point between the two curves quantifies the magnitude of the effect of the biocide. On a logarithmic scale, this parameter is called the log-difference or log-inactivation. This value is displayed in the legend for each timepoint. The log-reduction average and standard deviation are −1.48 and 0.03, respectively. These findings were consistent with the log-inactivation measured by conventional means^40^ (personal communication).

**Fig 4.**
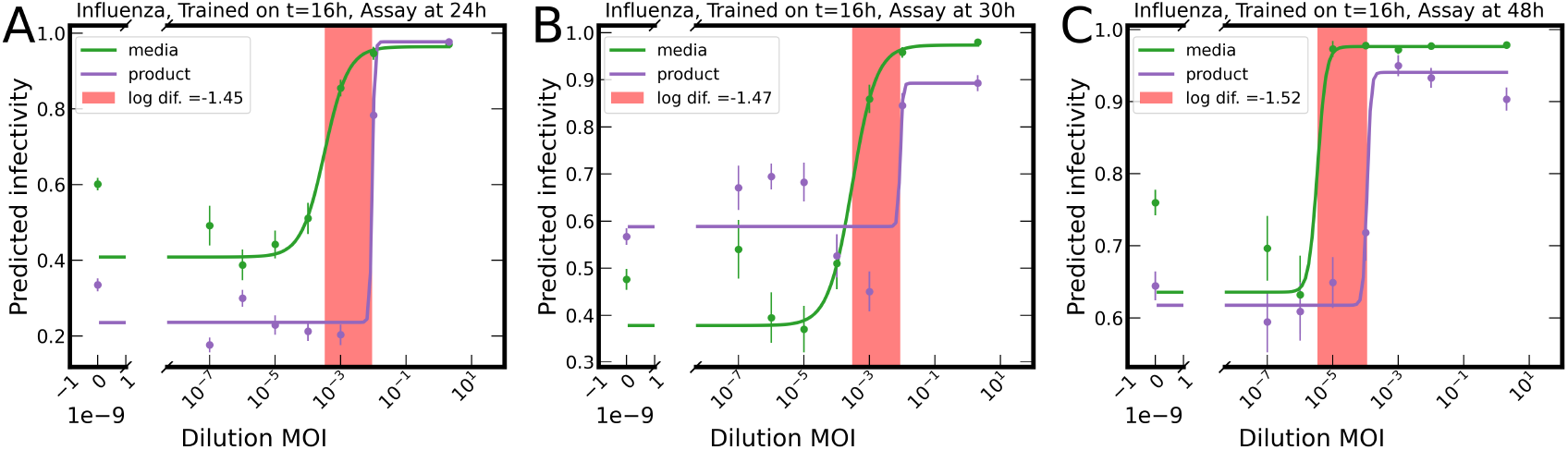
Inactivation assay on influenza A virus-infected cells treated with media or biocide. Infections were incubated for different times (24h, 30h, 48h; **A–C** respectively), imaged, and analyzed with a pre-trained AI (Influenza A at 16 hours post infection). X Axis: known virus dilutions; Y-axis: AI predictions of infectivity. Pink range represents the difference between IC50s for the fits to the media and product dilution series. The error bars show the standard error of the images for a given MOI.

It is noteworthy that the inactivation assay successfully quantified the log-inactivation at three incubation times well after the timepoint used to train the model. This demonstrates that the infectivity phenotypes present at 16 h were still being expressed in the cell culture at those later timepoints. The higher dilutions and longer timepoints allowed for multiple rounds of infection to be observed, expanding the range of log-inactivation observable in a single experiment.

Importantly, the log reduction measurement was stable across the three incubation times, with the standard deviation of 0.03 being approximately 2% of the mean. This provides further confidence for the robustness and precision of AVIA, and its applicability in real-world inactivation assays.

## Discussion

Incubation time occupies the vast majority of the total turnaround time of traditional assays for determination of infectious units. The timecourse analysis with AVIA revealed that infection phenotypes could be detected within hours after infection. This represents a dramatic reduction in the required incubation time for ML-assisted assays over traditional methods, in addition to the improvements in reproducibility, cost per sample, and throughput.

In HIV/TZMbl and Influenza/MDCK virus-cell line combinations tested, the model was able to distinguish infected from uninfected cells at the first measurement with relatively good accuracy. It is estimated that the time between infection and imaging the first time point would be between 15 and 45 minutes. This is likely to be well below the amount of time required for the infection to take hold and for the cells to show any early signs of virus production. One potential mechanism for these early stage phenotypes is cell signaling as a result of viral infection ^41,42^. We have not investigated the potential biological significance of these rapidly changing early phenotypes, but this could be of interest for further investigation. Artefactual shifts in phenotypes can be caused by CO_2_ and temperature shifts resulting from imaging plates outside on an incubator, and uneven evaporation during incubation. Differential effects from virus-containing media can be caused by cytokines or other factors carried over with the media at high-titer infections. These effects will be addressed in future studies addressing the mechanism of these early phenotypes.

Splitting images into tiles allows for a larger batch of training examples to fit into memory during each training step. Larger batch sizes expose the machine learning model to a greater variety of training examples simultaneously and decrease the volatility in each step of the training progression due to a larger sample size. However, a smaller area per tile means each positive tile is less likely to contain legitimate signals of infection. A greater number of smaller tiles also splits signals at tile boundaries and reduces contextual information. It has been argued that tiling images only serves to hurt performance ^43^. In this application, we believe there is more of a balance because a model trained on large tiles may perform better on the saturated infection in the training set, but we expect that the loss of ability to discriminate individual cells in large tiles and the reduced sampling will hurt performance in the sub-saturated infections used in real assays. The reduced number of training samples may also hurt the robustness of the trained models on unseen replicates. As expected, our preliminary observations are that larger tiles improve AI accuracy on the infected vs. uninfected cells (not shown). However this area deserves further investigation to determine the effect of reduced sampling, and of scoring large tiles that contain both infected and uninfected cells. It is also possible to train regressor models on intermediate dilutions of virus, so that AIs are exposed during training to the intermediate infection levels that they will be scoring when performing assays. Using large tiles with intermediate infections during training may overcome the effects of reduced sampling with larger tiles.

In most of the experiments described here, we could not detect differences between infected and uninfected cells by visual inspection, so the AIs appear to base their decisions on properties that a human observer cannot directly perceive. Some work has gone into trying to visualize what trained AIs perceive ^44,45^, including generative models that produce images to illustrate the differences they detect ^46,47^. We have not pursued trying to visualize or describe what our AIs use to differentiate infected from uninfected cells, primarily because our interest is currently limited to the utility of this tool in practical assays. We can however compare phenotypic similarities between cells infected with different viruses and time points (see Supplementary Materials sec. Timecourse Analysis for examples) and determine if these can be clustered by phenotype as perceived by an AI. We have used this technique previously to cluster phenotypes induced by RNA interference ^48^, age-related phenotypes in *C. elegans*^49^ and visual trends in modern art^50^.

As more viruses are characterized, we expect there to emerge clusters of similar phenotypes induced by viral infection in a certain host cell, which may correlate with infectivity pathways and antiviral defenses used by the different viruses. A map of infectious virus phenotypes may become useful when infectious viral load monitoring becomes more important in the context of control of seasonal and epidemic infectious events. This can be an important research tool for prediction of virulence, or potentially be used in a diagnostic setting such as environmental monitoring as an early indicator of the presence of infectious viruses.

The technique was validated on influenza A virus, HIV, MVA, poliovirus, vaccinia virus, adenovirus, and an adenovirus variant, spanning the four major types of viruses. On coronavirus 229E and MVA-infected cells, the trained models were robust across unseen biological replicates, without significant loss in accuracy. The consistency of this technique was demonstrated in an inactivation assay when the log-inactivation of an unknown biocide was measured at three different times after infection, achieving reproducibility within 2%.

We anticipate that this tool could be further developed into a broad and rapid diagnostic assay from infectivity phenotypes observed for a variety of viruses.

## Acknowledgements

This work was supported by NSF grant 2029707 to ViQi Inc. Support for JTK was provided by the Basic Science Core of the Texas Developmental Center for AIDS Research (AI161943). Support for TO, FS, AF, and EC was provided by the NCSU Comparative Medicine Institute, CMI, and the NCSU Center for Applied Virus Experimentation, CAVE. ProBioGen thanks Lina Hanisch and Nicole Seehase for expert technical assistance and Benjamin Werdelmann from Synentec for support in image acquisition with the Synentec Scientific device. ViQi and NCSU thank BioTek for their loan of a Cytation™ 1 to NCSU

## Author contributions

I.G.G. designed the study. T.O., F.S., A.F., E.C., N.H., I.J., and J.T.K. acquired the data. R.D., J.R.D., and I.G.G. wrote the code. R.D. and J.R.D. performed the analysis. I.G. supervised the work. All authors read and corrected the final manuscript.

## Methods

### Viral cultures

Datasets were prepared for 7 different virus/cell lines pairs from 6 different laboratories according to the training protocol described in Fig. 2 (mock-infected, synchronous high-MOI infection, and infections with 2-fold dilutions of virus). Except as noted in the text, infections and imaging were conducted on 96-well plates with flat plastic bottoms. Because the incubation time is low, the seeding density of the cells was relatively high (typically > 85% confluence), except as noted in the text. The dilution MOIs were predetermined in the different laboratories using TCID_50_ or plaque assay and assumed to be accurate to within a factor of 2.

The cultures for Corona 229E-infected Huh7 cells and Influenza A-infected MDCK cells were prepared as described in et al.^51,^ and Crisci et al.^52^, respectively. The 229E images used in Supplementary Fig. 1 were captured at 20x and 40x using a manual microscope. Other 229E and influenza/MDCK images (see Supplementary Fig. 9) were captured on a Cytation 1^37^ at 20x magnification. Modified vaccinia ankara (MVA) wildtype (WT) and CR-infected CR.pIX cells^53^ were prepared as described in Jordan et al.^54^, and the images were captured with a Synentec NyONE^36^ at 20x. Human immunodeficiency virus (HIV)-infected TZMbl cells were prepared as described by Wang et al.^55^ and the images captured with a Cytation 5^38^ at 20x.

Adenovirus-infected HeLa cells were prepared as described in Jogler et al.^56^, vaccinia-infected BHK-21 cells were prepared as described in Earl et al.^57^, polio-infected HeLa cells were prepared as described in et al.^58^. Influenza-infected MDCK cells (see Supplementary Fig. 9) were prepared as described by Kim et al.^59^ and the images were captured with a Synentec CellaVista^35^ at 20x.

### Imaging

Each well on the 96-well plates was imaged 13-25 times using a grid of non-overlapping images. Images were collected every 2-4 hours before the onset of visible CPEs. Imaging was done with a high-throughput microplate imager or manual microscope (ZEISS Axio Imager/AxioCam HRc) as noted above. The sensor sizes were as noted in Table [2; virus table]. The focal plane of the images was set to the interior of the cells as opposed to the tops of the cells as is commonly done for brightfield to optimize cell segmentation tasks. The images were stored as 16-bit grayscale TIF-files or 48-bit RGB (16-bit per channel), in which case they were converted to grayscale using the NTSC formula^60^.

### AI training

The classification model is a convolutional neural network (CNN), which uses a series of convolution kernels to extract features from input images at increasing levels of abstraction ^61^. This model was implemented using Tensorflow^62^. The weights of these kernels are initialized to those pretrained on ImageNet^63^. Bootstrapping on the knowledge acquired from those models allowed the AVIA models to converge faster and at a higher accuracy ^64^. After each cycle through the full training data (an epoch) the data is transformed with flips and rotations to increase the scope of the training examples.

Depending on the strength of the morphological signals from the infection, and the allowed time budget for training, models of varying complexity are trained. A single CNN based on EfficientNetB0^65^ can be used as a rapid model with modest accuracy as a stand-alone model for training. On the other end of the scale, a total of sixteen CNNs can be trained with different base models and learning parameters (hyperparameters), and their predictions fed into an ensemble classifier such as a support vector machine (SVM) or random forest classifier ^66,67^. In our experiments, three other CNN architectures were used including MobileNet^68^, InceptionResNetV2^69^, and VGG^70^; and hyperparameters with two options for dropout^71^; and two options for learning rate. Future work will also include tiles with multiple sizes.

In initial experiments, cells were seeded at a lower confluence more typical of conventional infectivity assays that have long incubation times to allow for cell growth. Because of the short incubation time for AVIA, in subsequent experiments, cells were seeded at higher density or > 85% confluence to maximize the number of cells and thus the amount of information available to the AI when the cells are imaged. In the earlier, more sparsely seeded datasets, we trained a prefilter network to eliminate tiles that did not have sufficient cellular material. The image data for this network was the same as for the infectivity network, except tiles were manually labeled that had little to no cellular material so that they would be eliminated by the prefilter. This prefilter network was easily trained to approximately 92% accuracy on each occasion (not shown).

### Ensemble AI

In ensemble models, numerical outputs of multiple CNN models (marginal probabilities of a given tile being infected) are fed as input features into an automated feature classifier trainer developed by ViQi. This trainer uses scikit-learn^72^ to try several different feature normalization, scoring and selection techniques coupled to several different high-performing classifiers. The trainer optimizes appropriate parameters for the algorithms at each stage and automatically selects the best-performing model^73^ and corresponding parameter set.

## Supplementary Material

### Training optimization

There are many free parameters in designing the experimental setup and the model hyperparameters that must be investigated and tuned in order to give an AI model the best chance of finding features to discriminate infected from uninfected cells effectively. In this section, we discuss several investigations and their effect on the parameter choices. These experiments were performed on several datasets with different virus/cell line pairs. This way, the findings are not bound to just one combination of cell line and virus. In some cases, the experiments are presented on multiple datasets to provide further confidence in the findings or to illustrate differences between datasets. In other cases, it was only possible to perform the experiment on one dataset.

### Effects of magnification

With modern cameras (1k X 1k pixels or greater) and optics, 40x objective lenses typically produce images at the resolution limit expected for light microscopy without substantial oversampling, or ∼ 0.25μm/pixel with 2k x 2k cameras^1^. However, 20x objectives, especially with 2k or 4k cameras, provide resolution near the resolution limit for visible light with several practical benefits over 40x. Common plastic-bottomed tissue-culture plates are routinely used with 20x objectives rather than more expensive glass-bottom or other specialized plates. The focal plane is typically thicker with 20x objectives, allowing us to image more of the intracellular volume. Lastly, with 20x objectives, the amount of cellular material per image is increased by a factor of 4 relative to 40x, or the number of images per assay could be reduced by a factor of 4, which would be a substantial savings in imaging, upload and processing time.

For example, training data for Coronavirus 229E-infected Huh7 cells was plated for two replicates – one was captured at 20x magnification and one was captured at 40x magnification. The same microscope and camera sensor was used to capture the same number of images for both sets. The 20x dataset therefore contained more cell examples for the model to learn from, but less detail per cell. Both of these two datasets were resampled to create a total of four datasets – the 40x raw data was downsampled by a factor of two to match the resolution of the 20x raw data, and the 20x raw data was upsampled to match the resolution of the 40x data. The downsampled 40x dataset contained the least amount of tiles of the four datasets (4x less than the raw datasets) and the 20x upsampled dataset contained the most amount of tiles (4x more than the raw datasets). To control for the variable number of tiles, one experiment was run with the number of tiles in each dataset limited to the number in the smallest set (40x downsampled; Supplementary Fig. 1A), and a second experiment was run with the uneven number of tiles (Supplementary Fig. 1B). A fresh model was trained for each of the eight datasets and the accuracy was evaluated on test images not previously seen by the trained models.

When the number of tiles is held constant, if the amount of cellular content is the primary driving factor, as opposed to high-resolution cellular features, the model should perform approximately equally well on 40x raw and 20x upsampled, and similarly on 20x raw and 40x downsampled. Instead, we see that the 20x dataset performs best at each resolution, indicating that the reduced depth of field in the 40x data, and therefore the consistency in the focus across the image, places a significant limitation on the model performance. This discrepancy between 20x and 40x magnification becomes even more pronounced as the increase in the number of tiles for 20x over 40x is allowed to skew model performance. These results, in addition to the practical benefits for image acquisition, provide a strong indication that 20x magnification is more of an optimal condition for AVIA experimental setups. Of course, the relevant morphological features will be different for different viruses, and high-resolution features may still become a deciding factor in some viral infections.

**Fig. 1.**
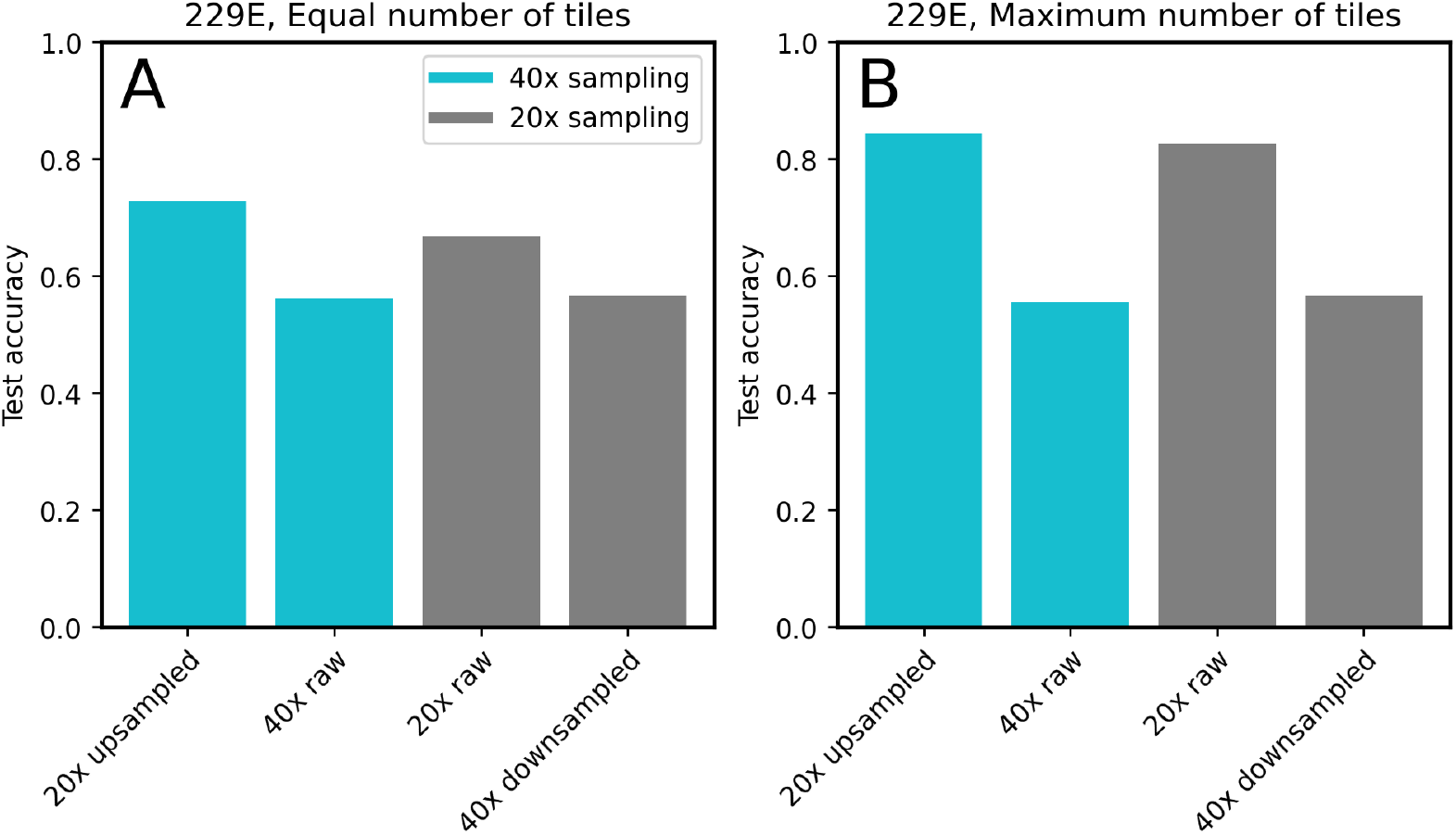
Exploring the effects of magnification on training accuracy. Raw image data at 20x and 40x are from two different replicates and magnifications of Coronavirus 229E infection of Huh7 cells. Blue bars are at the spatial sampling of the raw 40x data, and gray bars are at the spatial sampling of the raw 20x data. **A**: the number of tiles was held constant at the minimum of the four experiments (40x downsampled). **B**: the maximum number of tiles available were used for each dataset.

### Timecourse analysis

Training a multi-class model with a positive class and negative class for each timepoint reveals the phenotypic similarity between all classes simultaneously (see Supplementary Fig. 2A). Each row represents a different known class (e.g. row 1 is uninfected cells at 0 hours post-infection). The raw AI output for a given image tile is the set of marginal probabilities for all classes defined in the classification run, and the selected class prediction is the one with the highest value. Thus, the column values in each row are the distribution of class predictions for a given row’s true class. The correct predictions are on the upper-left to lower-right diagonal in this view, and the remaining values indicate classes confused by the AI with the correct class. For example, a comparison can be made between the negative classes at all timepoints (top left quadrant). This reveals the kinetics of cell growth in the absence of infection. Or we can compare the confusion between the positive classes at all timepoints (bottom right quadrant), which reveals the kinetics of the infection convoluted with any residual cell growth. We can also compare positive and negative classes at a single timepoint to estimate what a binary model using only that single timepoint would achieve. The set of four points in a binary model would mimic the standard binary confusion matrix, with the caveat that the multi-timepoint model is also trained on the other timepoints while a binary model would be trained on a single timepoint (or possibly two adjacent ones).

As time progresses in the uninfected cells (going down the top left quadrant of Supplementary Fig. 2A), the distribution around the true positive diagonal broadens, indicating that there is a larger amount of confusion between timepoints, and the cells are becoming less distinct with time. A similar trend occurs in the infected population. These trends are relatively smooth, indicating gradual, monotonic progressions of the cell morphology with time.

There is a relatively strong true positive and true negative signal at 0h after infection as indicated by the relatively large signal (0h+, 0h+), counter to the intuition that the cells will have had insufficient time to display signs of infection this early after infection. This indicates that a different mechanism is at play for this timepoint, which might not correlate well with infectivity and requires further investigation. We discuss possible explanations for this in the discussion.

The 24h timepoint has the highest accuracy among those on the true positive diagonal that can be considered an infectivity signal. Therefore, for HIV, 24h appears to be the optimal time to train a binary model at and use for assays. In this case combining data from adjacent timepoints should help training accuracy as the neighboring cell phenotypes appear similar.

The changes associated with uninfected cells can be removed from the infected cell timecourse leaving only the kinetics associated with infection. A way of doing this, that assumes the phenotypic changes in the infected culture are superimposed on the phenotypic changes in the uninfected culture, is to perform a vector subtraction of the uninfected probability distribution from the infected one at each timepoint. Euclidean distance between infected probabilities at each timepoint can then be used to calculate phenotypic distances to visualize with dendrograms.

Supplementary Fig. 2B shows the dendrogram for the HIV infection through time. The phenotypic distance between any two classes in the dendrogram is reflected by the total distance along the branches that connect them. Up to the third timepoint, the progression is fast based on the large branch length between 0h–3h, and 3h–6h. Between 6 h and 18 h the infection undergoes a relatively slow transition and the progression accelerates at the later timepoints. The classes getting progressively further from 0h validates that the infection is monotonically progressing with time. It is important to point out that the order of the classes was never presented to the AI in training, so its ability to reconstruct the order of the timepoints indicates that it is based on progressive phenotypic changes in the cells.

**Fig. 2.**
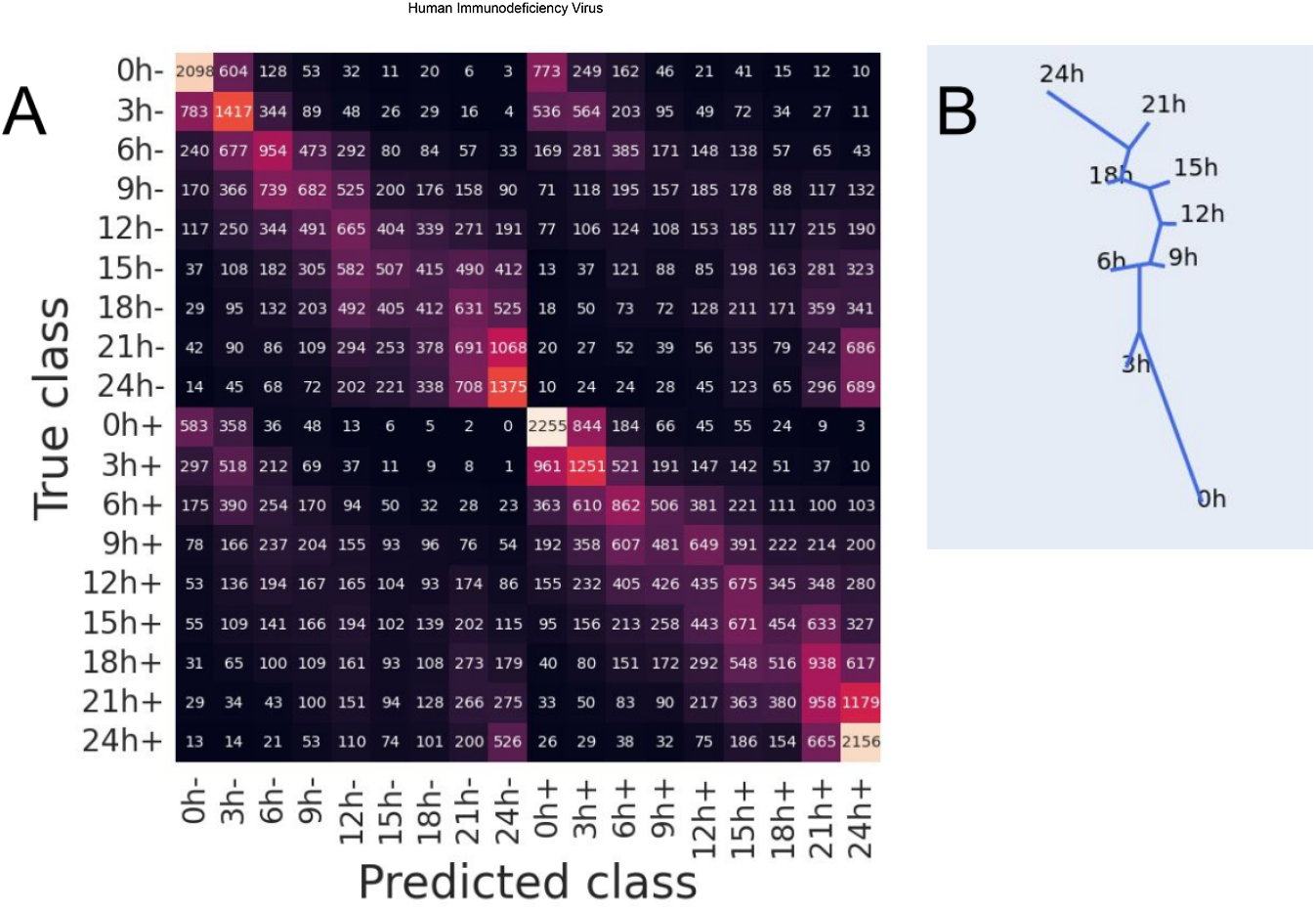
Simultaneous time and infection multiclass models for HIV displayed in confusion matrix and dendrogram formats. In **A** the class number represents the number of hours after infection and symbol represents the infection condition: infected (+) and uninfected (-) cells. Rows denote the distribution of the model predictions for each class. The correct calls for each class are on a top-left to bottom-right diagonal, and any signal outside of the diagonal represents confusion between the known class in the row and the predicted class in each column. **B**, the phenotypic progression of the infection after removing uninfected cell classes. For a description of how these are interpreted as phenotype similarities between classes, see text.

The timecourse for influenza A-infected MDCK cells is shown in Supplementary Fig. 3. There is a large amount of confusion shown among the uninfected cells with a phenotype transition happening between 4h and 6h. The infected population shows little confusion at time: 2h, 4h and 10h after infection, and a significant amount of confusion at 6h and 8h.

Supplementary Fig. 3C compares multiclass timecourses for influenza A-infected MDCK cells analyzed using two types of machine learning techniques: CNNs as done throughout this study, and a feature-based classifier. In this case, a 10x objective was used for imaging, and the same images with similar tile sizes were analyzed by the two different machine-learning techniques. The predictions from the feature classifier used predetermined image analysis features (CHARM features^2^) such as textures and polynomial decompositions, and the same auto-ML software described in Methods. The predictions from the CNN model (the algorithm core of AVIA) used features that it fabricated for the specific training set. Interestingly, the same phenotypic patterns across time can be seen in the feature classifier confusion matrix albeit with a lower signal to noise ratio. The fact that a measurement as orthogonal as a feature classifier based machine learning algorithm versus a CNN algorithm displays similar patterns in the confusion matrix illustrates that these patterns can not be an artifact of the machine learning algorithm itself.

**Fig. 3.**
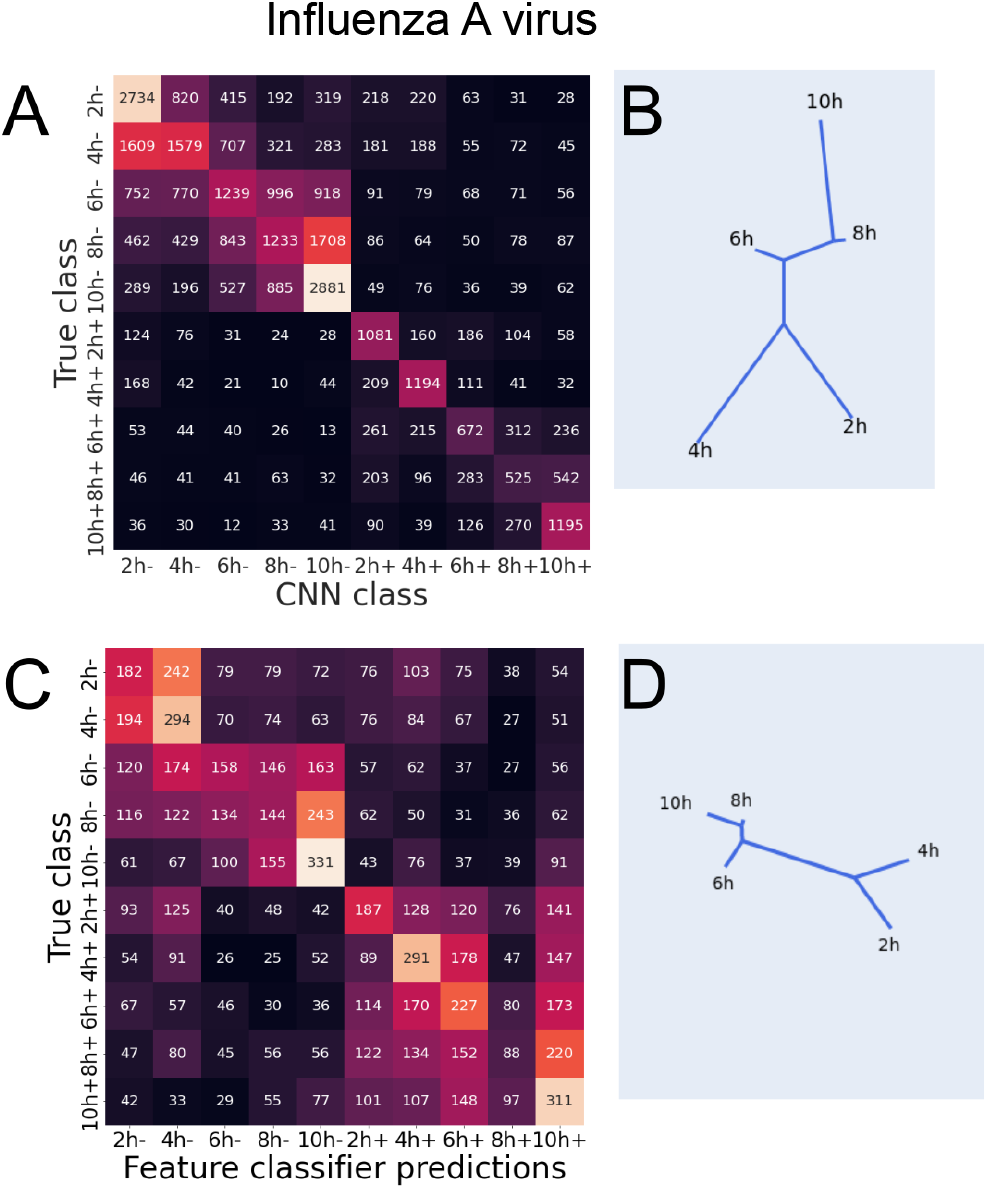
Comparison of two types of machine learning methods on time and infection multiclass models for influenza A. Format for the confusion matrices and dendrograms are the same as Supplementary Fig. 2. **A** and **B**, the predicted dynamics according to the CNN-based model used in AVIA. **C** and **D**, the predicted dynamics according to a feature classifier-based model.

## Robustness to experimental conditions

The central challenge in machine learning engineering is developing a robust algorithm that generalizes to new input data from a limited amount of training data and resources. In this section, we evaluate the generalizability of models across experiments. The current AVIA workflow requires new training data and algorithms for each new virus, cell line, and laboratory, which corresponds to its practical application. Assessing the performance of a model trained across those domains may be of interest for diagnostic purposes, or potentially to attempt to discover phenotypic classes of infection common to multiple viruses.

### Replicate robustness

**Fig. 4.**
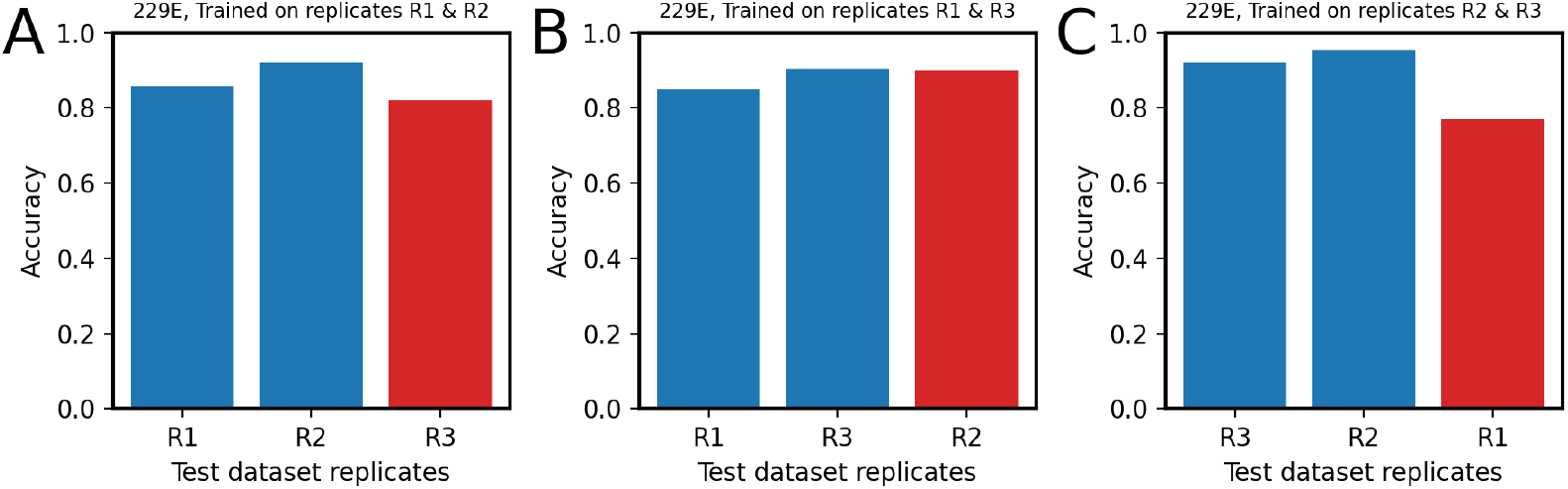
Comparing extrapolation of models trained on two biological replicates to a third replicate. Panels **A–C** display the results for replicate pairs (R1,R2), (R1,R3), and (R2,R3), respectively. The blue bars represent the replicates that the model was trained on and the red bars represent test data from a replicate not used in training.

Collecting results from several biological replicates accounts for variability in cell cultures. Collecting 3 biological replicates allows for a model to be trained on data from 2 replicates, so that the model is not overtrained on any one replicate. The third replicate can then be used to check the model’s generality. This experiment was conducted on replicates of Coronavirus 229E-infected Huh7 cells. This dataset had low seeding density (∼50%) so the prefilter network (see Methods) was applied to all images. Supplementary Fig. 4 shows the performance of these 3 models on unseen images from the two replicates used for training (blue) and images from the remaining replicate (red). Test images from each replicate were withheld from the models during training and were used for the assessments. If any of the models suffered from severe overtraining, the accuracy on the unseen replicate would be much lower than the two replicates used in training. In each case, the performance on the unseen replicate is comparable to that of the training replicate. For example, in panel C the test replicate accuracy is still well above the equal probability binary classifier noise floor of 50%, and in panel B the test replicate accuracy actually exceeds one of the train replicates. This was an important demonstration that the AIs are not overly sensitive to specific cell growth conditions or infections.

### Temporal robustness

Supplementary Fig. 2 showed that multitime classifiers have some confusion among nearby timepoints. This section frames the question in the context of a series of binary classifiers, which allows for a direct test, free of contamination by the other timepoints, and more closely mimics the training of AIs for subsequent titration assays. For example, Supplementary Fig. 5 shows the performance of 4 models trained on HIV-infected TZMbl cells at times 0, 9, 15, and 24 h after infection, and 4 models trained on Coronavirus 229E-infected Huh7 cells at 1, 4, 8, and 12 h after infection – well before CPEs are visible^3^. For the HIV experiment, the test images were from wells randomly distributed across either class, and for the 229E experiment, testing was done on a biological replicate left out of training. The same train/test partition was used for each timepoint.

The cells at each timepoint will be at a different stage in their progression, so it is expected that the model accuracy would be substantially lower at the other timepoints. This form of overtraining explains the height difference between the blue and red curves. However, the heights of the red curves are well above the binary classifier noise floor of 50%, indicating a modest degree of robustness across these time differences.

With the models trained on HIV (Supplementary Fig. 5 A–D), the model constructed with 0h (panel A) training data has the highest accuracy on test data from the same timepoint (blue) among the four models, as well as the largest difference between that value and the mean of the data from the other timepoints (red). This provides further evidence of an early phenotype that degrades at later timepoints (as first noted in Supplementary Fig. 2).

**Fig. 5.**
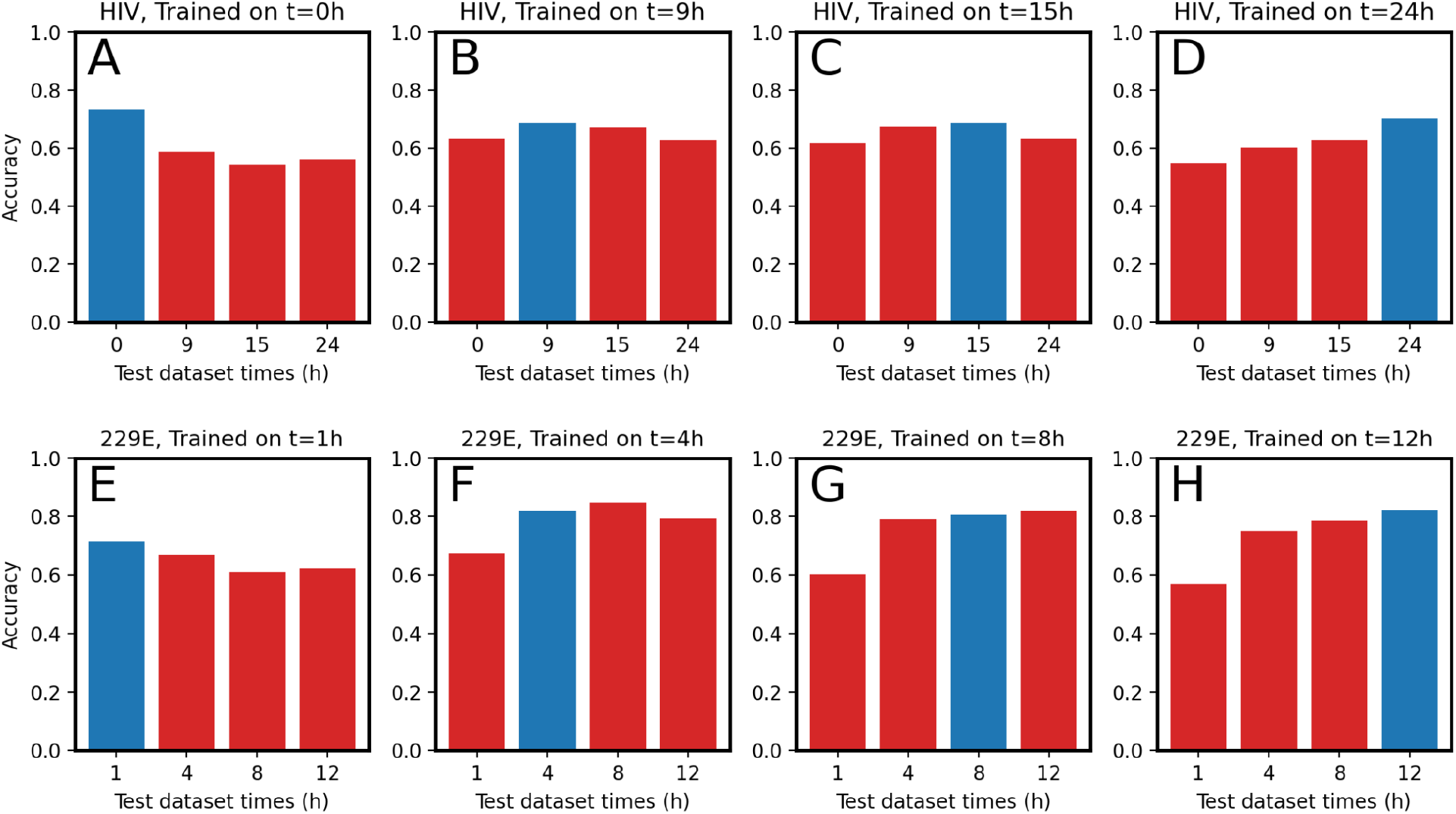
Comparing extrapolation of models trained at one time after infection to other timepoints. Panels **A–D** display the results for models trained on HIV infections at 0, 9, 15, and 24 h post-infection, respectively. Panels **E–F** display results of models trained on Coronavirus 229E infections at 1, 4, 8, and 12 h post-infection. The blue bars represent performance on test images from the timepoint that the model was trained on and the red bars represent test data from the other timepoints.

In Supplementary Fig. 5 panels E–H, models trained 1h post-infection (panel E) perform adequately on the 1h timepoint, but perform slightly worse on subsequent timepoints. Models trained at 4, 8, and 12 h post-infection (panels F–H) all perform well on images from 4, 8, and 12 h post-infection, but comparatively worse on images from 1 h post-infection. Similar to the HIV 4 model experiment, there appears to be an early short-lived phenotype, but over the later 3 timepoints, the second phenotype is more persistent.

### Strain robustness

Different viral strains can yield radically different characteristics of infection in cells. Without imaging in fluorescence, we can ask if an AI trained on one variant can accurately distinguish cells infected with the other variant. This is important for validating that AIs trained on one dataset are not bound to the same set and can extrapolate to different infections, potentially with different related viruses on the same cell line. In this experiment, CR.pIX cells were infected with two variants of MVA, wildtype (WT) and strain CR, that express different fluorescent proteins. Ensemble models consisting of 16 base CNN models were trained on the CR (Supplementary Fig. 6A) or WT variant (Supplementary Fig. 6B) of MVA, and were used to evaluate images from the other variant. The 3 sets of images used to evaluate models were those used internally during training (blue), images from the same variant withheld from training, and images from the other variant (red). The performance on the previously unseen strains is substantially greater than random indicating that the AI can successfully generalize to the unseen strain. Although the accuracy of the predictor dropped significantly when evaluating the other strain, it remained substantially above noise. This is a good indication that models may be able to distinguish these virus infections, but also report a measurable phenotypic similarity.

**Fig. 6.**
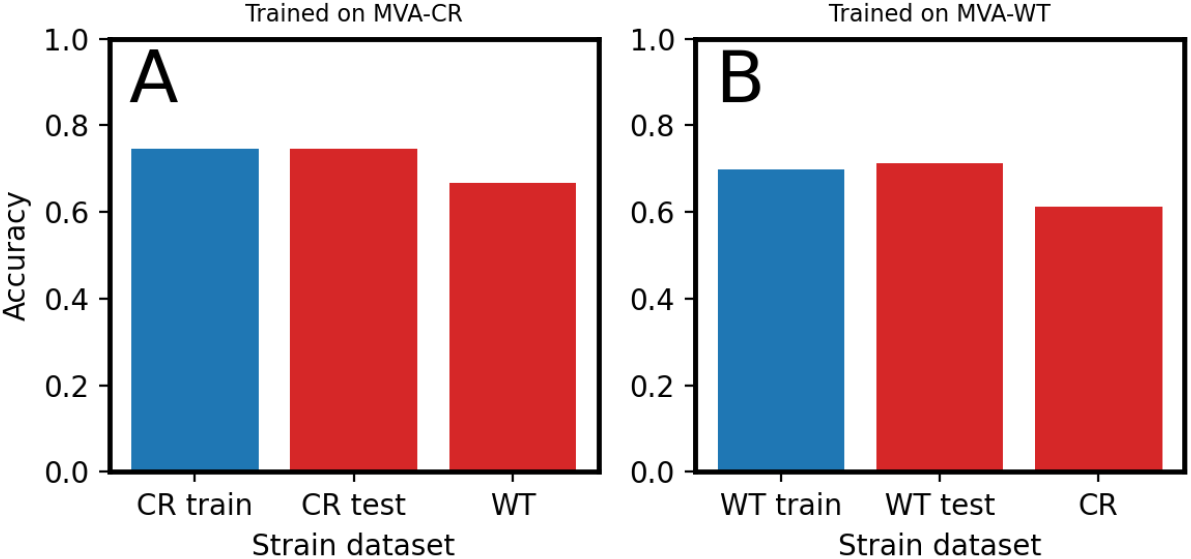
Comparing extrapolation of a model trained on the CR strain of MVA to another strain. Panel **A** displays the performances of the model trained on the CR strain of MVA and panel **B** displays the performances of the model trained on the WT strain. The blue bars represent the data that the model was trained on and the red bars represent test data.

This experiment can be extended by applying these two models to validation protocol data from both CR and WT strains (Supplementary Fig. 7). Despite the drop in performance across strains, the correct sigmoidal trend with MOI is achieved and the linear range is consistently within the range of measured dilution MOIs. The linear range is narrower by more than 40% for the unseen strain in both instances, encompassing fewer of the dilution MOIs, which is in line with the lower performance. The performance of the CR model predicting WT (Supplementary Fig. 7B) fits well with the expectation that the CR strain escapes the cells more easily, often causing much weaker CPE than WT. In Supplementary Fig. 7C, an AI trained on the weak CPEs of CR can predict the stronger WT CPEs (bottom row), but an AI trained on the stronger WT CPEs is not able to score the weaker CR CPEs (top row). A model could potentially be trained with both variants to desensitize it to the differences between them and create a calibration curve with a higher signal-to-noise ratio.

**Fig. 7.**
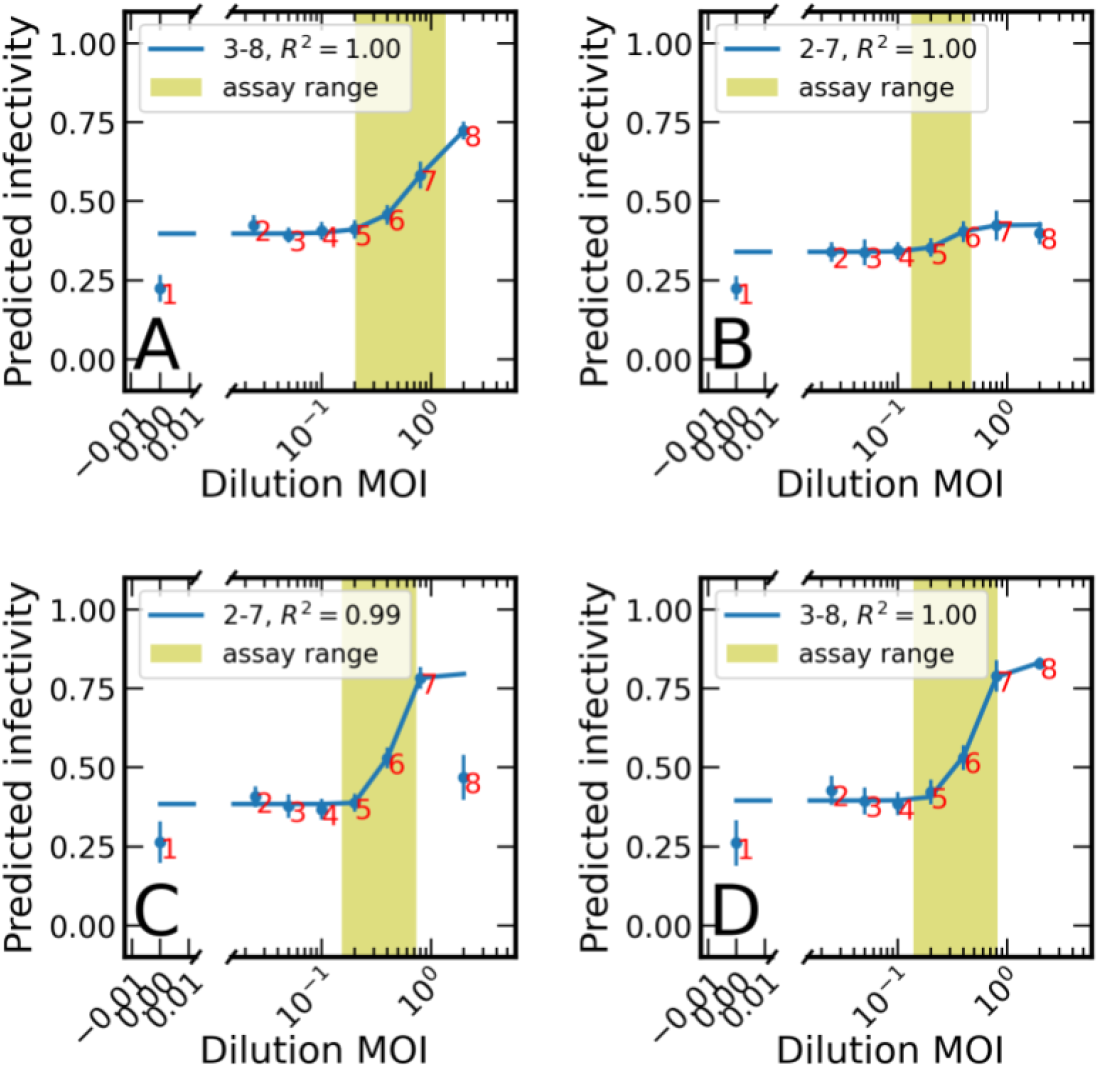
Comparison of the dilution calibration curves for models trained on two different MVA strains. **A**, the result of training and validating on the WT strain. **B**, the result of training on the WT strain but predicting on the CR strain. **C**, the result of training on the CR strain but validating on the WT strain. **D**, the result of training and validating on the CR strain. The fit with the highest R^2^ as well as the ranges of datapoints used in that fit is shown in the legend. The error bars show the standard error of the images for a given MOI.

### Laboratory strain and temporal robustness

The larger the number of differences between two datasets, the less likelihood there is for a model exposed to only one of those datasets to achieve generality in discerning infectivity features in the other dataset. For example, in Supplementary Fig. 8 we compare the similarity of two different datasets of the same virus from different laboratories. The dataset from laboratory A contains influenza A virus-infected MDCK cells, at 10h after infection, on a BioTek Cytation 1. This instrument used a 10x objective and a 1k camera for a resolution of 0.67 μm/pixel. The dataset from laboratory B contains the same virus strain and cell line, but at 16h after infection and on a Synentec CellaVista. This instrument used a 20x objective and a 2k camera to obtain images at 0.65 μm/pixel. Despite the different magnifications and cameras, the images were not further adjusted due to the close match in resolution. In this experiment, the accuracy achieved on the unseen data is significantly less than the test data from the same laboratory. The accuracy of the laboratory A model on the laboratory B data is marginally better than the noise floor indicating that perhaps there may be some common signals shared between the datasets, but the noise associated with the different samples and their conditions is overwhelming the signals. It appears as though in order to generalize to samples from unseen laboratories, data from several will have to be combined to desensitize the model to these differences. This is left for future work.

**Fig. 8.**
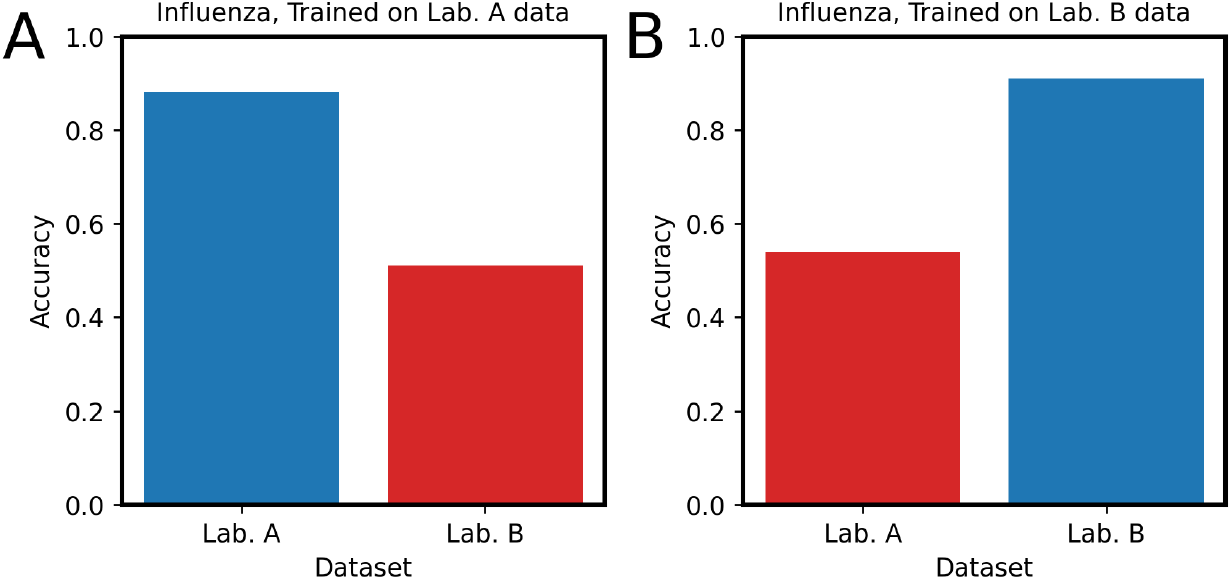
Comparing extrapolation of a model trained on Influenza A virus to a different laboratory, device and time after infection. Panel **A** displays the performances of the model trained on laboratory A influenza experiment and panel **B** displays the performances of the model trained on the laboratory B influenza experiment. For more detailed description, see text. The blue bars represent the data that the model was trained on and the red bars represent test data.

### Comparison to GFP

A direct comparison between AVIA and FFA was conducted using CR.pIX cells infected with modified vaccinia ankara expressing green fluorescent protein (MVA-CR). Several models were trained on the MVA-CR images using a range of hyperparameter settings. It was found that, for this dataset, an EfficientNet-based model with 40% dropout had the highest performance on training data – we therefore use that model for the investigations in this section. It was observed that the AIs can detect differences between infected and uninfected cultures at 4 hours.

In this section, we define a criteria for tiles containing GFP to compare them to tiles predicted as infected by AIs. A multi-resolution filter was used on the GFP channel to eliminate low-frequency noise. Then a global threshold was used to identify pixels with GFP signal. Tiles containing any GFP pixels were counted as infected. GFP infectivity for an image was the sum of GFP-positive tiles divided by the total number of tiles, same as for AVIA infectivity. The means and standard error for AVIA and GFP infectivities for images at each MOI were used in Supplementary Fig. 9. The colors refer to three sources of data. The purple datapoints were taken from a plate containing only mock and synchronous MOIs used for training, and the remaining colors refer to different rows from a plate containing dilution MOIs for validation. The orange datapoints were from rows at the edge of the plate, which may suffer from plate edge effects. This set of dilutions was also missing images from the 0.8 MOI dilution. This set of dilutions is displayed, but was not used in the fit. The correlation (Pearson r = 0.87) shows that infectivity tracked by AVIA in brightfield images is well correlated with infectivity tracked by GFP in the corresponding fluorescence images. It also illustrates the potential consequences of plate edge effects, as well as the excellent reproducibility between replicates when plate edge effects are factored out.

**Fig. 9.**
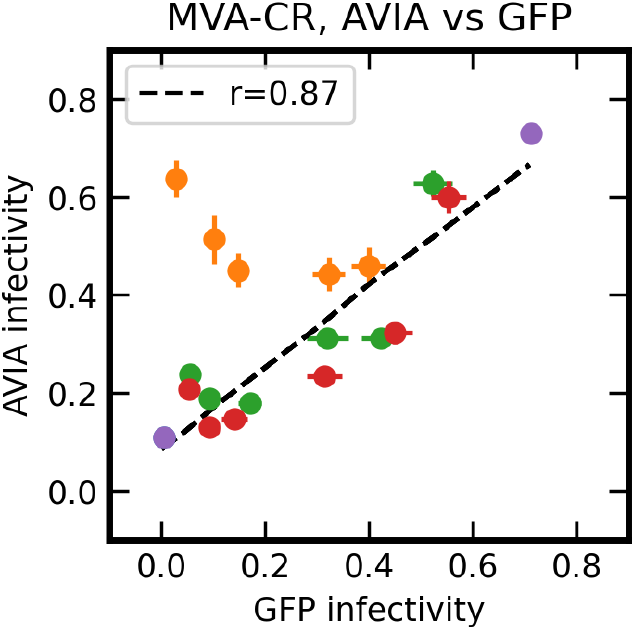
A comparison of the average infectivity predicted by GFP to the predicted AVIA infectivity for each dilution on MVA-CR-infected cells. The error bar displays the standard error of the distribution of predictions for MOI under investigation. The legend displays the correlation coefficient between the two methods using only the red, green and purple replicates.

**Fig. 10.**
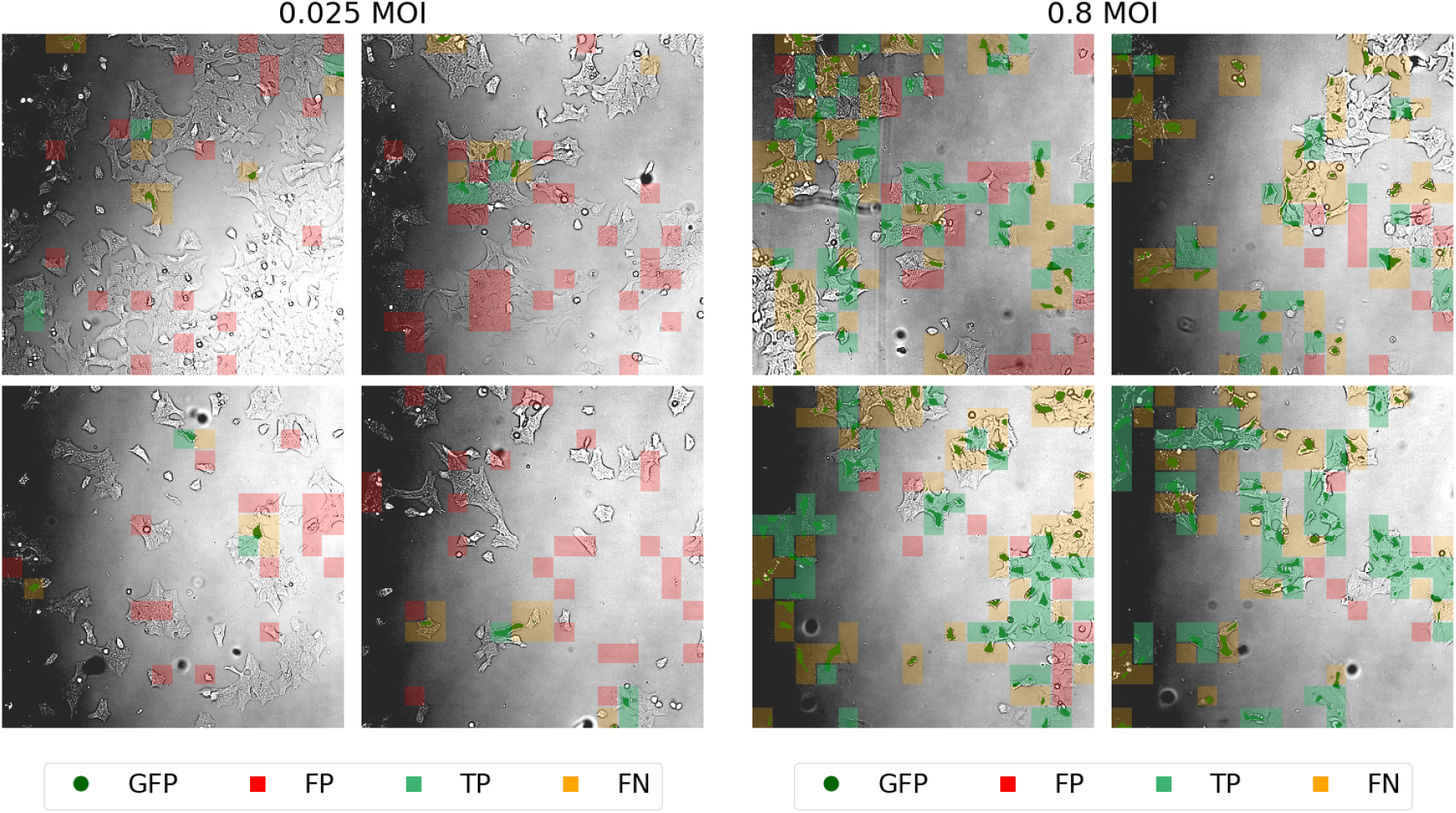
Cell infection validation on MVA-CR. The left 2×2 matrix contains brightfield images at the highest virus dilution and the right 2×2 matrix contains brightfield images at the lowest virus dilution. In each image the GFP channel is overlaid on top of the brightfield channel, along with the color-coded AVIA tile predictions with false positives (FP) in red, true positives (TP) in green, and false negatives (FN) in orange. True negative tiles are not displayed.

Supplementary Fig. 10 shows correlation between GFP-infected and AVIA-predicted tiles. The uneven illumination within and between brightfield images does not appear to affect AVIA predictions. The 0.025 MOI images show a relatively high false positive rate at low MOIs, including several false positives at locations that don’t contain any cells. This indicates that the false positive rate could be helped by training a more thorough background prefilter network. Interestingly, while we have shown that AVIA infectivity scales with GFP infectivity when performing a mean of each well, on an image to image basis, we found the locations of AVIA and GFP do not always align. This indicates that AVIA may be using a different signal in different parts of the cells to those that fluoresce in GFP. This effect deserves further investigation in future work.

## Example Training Report

Viral Infectivity Assay Al Training Report on Adenovirus ViQi, Inc.

## Abstract

ViQi, Inc. has developed an Al model for rapid and automated viral titer quantification. We applied this model to HeLa cells infected with Adenovirus at time 24 hours after infection. The images were captured with a Synentec CellaVista [1] microplate imager by Weyland-Yutani Corporation. On training data, the model was able to discern infected from non-infected tiles with 80% accuracy on unseen data. The dilution assay compares dilutions of the virus stock with known titer with predictions by the trained Al. A fit to a sigmoidal function has an *R*^2^ of 1, and an assay sensitivity range between 0.053 and 0.61 MOI.

### 1 Input Data Statistics

The input data can be separated into two types: the training protocol and the dilution assay. The training protocol contains images at Multiplicity Of Infection (MOI) 0.0 for the non-infected class and images at MOI 2.0 for the infected class (at which point ∼86% of cells should be infected, based on Poisson statistics). The dilution assay consists of images taken of a 2-fold dilution series to determine the linear assay range of the trained Al. Figure 1 summarizes the image statistics for the two types of protocols. Example images at several MOIs can be found in the Appendix.

**Figure 1:**
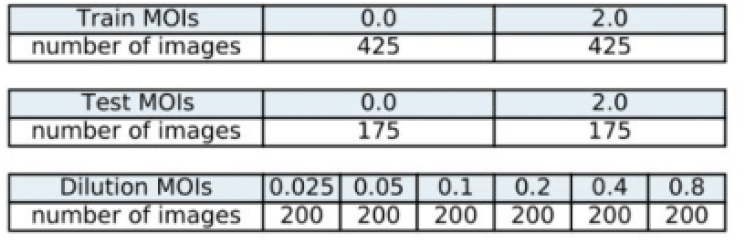
The number of images in the train, test and dilution assay wells separated according to dilution MOI. The Al learns from the train dataset, it’s learning performance is evaluated with the test dataset, and it’s performance as a quantitative infectivity assay is evaluated with the dilution assay dataset.

**Figure 2:**
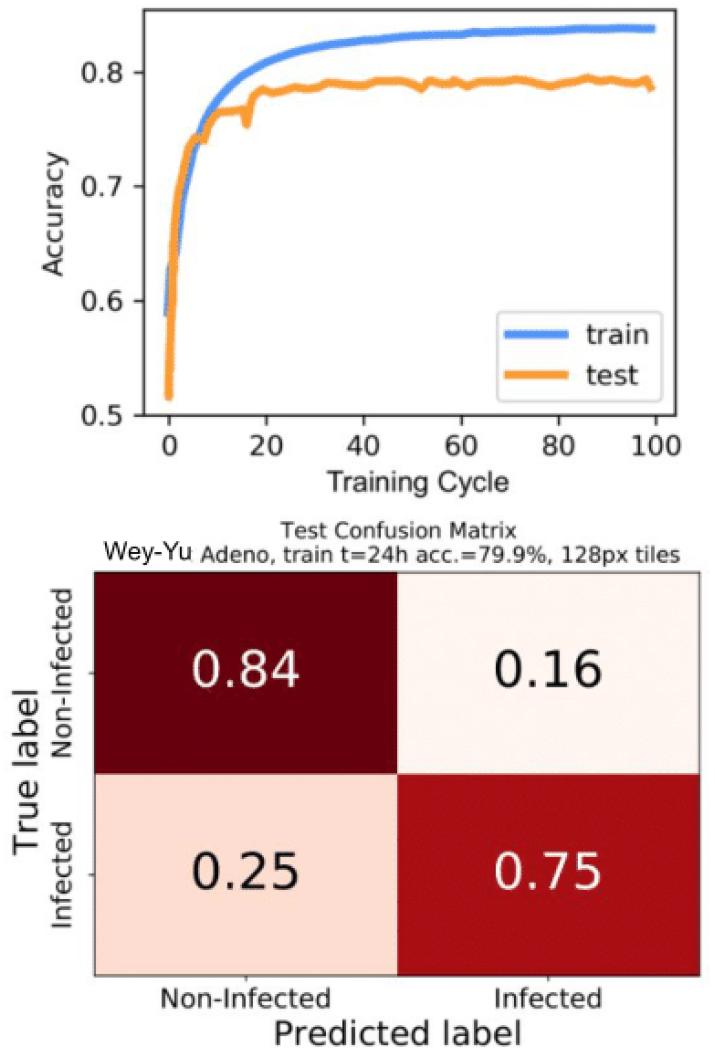
(top) the binary accuracy of the Al model on train and test (unseen) data as it trains. The fact that the training plateau is above the testing plateau indicates that further improvements could be made by introducing a greater scope of training data to the Al or fine tuning the parameters more, (bottom) the performances of the Al for both classes on the test data. This matrix shows that the Al tends to confuse the infected class more but a different threshold could counteract this if neccessary.

### 2 Model Quality

The infectivity Al was constructed by training a convolution neural network and transfer learning from ImageNet [2], After each cycle through the full training data (an epoch) the data was transformed with flips and rotations to increase the scope of the training examples. The learning curve for the Al as a function of epoch is shown in Figure 2 (top). The fact that the training plateau is above the testing plateau indicates that further improvements could be made by introducing a greater scope of training data to the Al or fine tuning the parameters more. The test data peak accuracy of 80% is higher than 52% of the models we’ve trained across all virus, conditions, and time points. The Table in Figure 2 (bottom) is a confusion matrix showing the performances of the Al for both classes on the test data. This matrix shows that the Al tends to confuse the infected class more but a different threshold could counteract this if neccessary.

### 3 Dilution Assay Results

The predicted infectivity was measured for each image in the dilution assay. This was done by counting the number of tiles with infectivity above the threshold 0.5. These scores were averaged for each MOI and plotted on the Y axis in Figure 3. with the X axis set by the known MOI from the dilution series. This curve was then fit with the four parameter logistic model (4 PL)

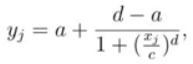

where *y*_*j*_, is the infectivity at MOI *x*_*j*_,*a* is the infectivity of the upper asymptote, *d* is the infectivity of the lower asymptote, *c* is the MOI at the inflection point of the curve, and *b* is related to the slope at the inflection point [3].

In order to better identify the upper and lower asymptotes, a series of fits were performed after cummulatively masking points starting from the high and low MOIs and then evaluating the *R*^2^ on the remaining datapoints until the minimum valid number of datapoints (5) was reached. The test datapoints from the training protocol at MOI 0.0 and 2.0 were included in these fits labeled as points 1 and point 8 The three fits with the highest *R*^2^ as well as the ranges of datapoints used in those fits are shown in Figure 3. The *R*^2^ for the best performing data range is ∼ 1 indicating a high goodness of fit.

The range of MOIs where an assay could be conducted, or the linear region, was determined from the intercept of the linear function extending from the inflection point and the upper and lower asymptotes. The effective assay range was determined to lie between 0.053 and 0.61 MOI. This corresponds to a 12 fold range, which is comparable to that used in a traditional plaque assay (with a countable range of 10-100 plaques per well) and much wider than those used in TCID_50_ (typically 2-fold) [4], To cover a broad range of possible titers, 5 fold dilutions within this range is recommended for the assay The titer of the undiluted virus can be determined from the points within the linear range, the dilution, and the cell plating density.

**Figure 3:**
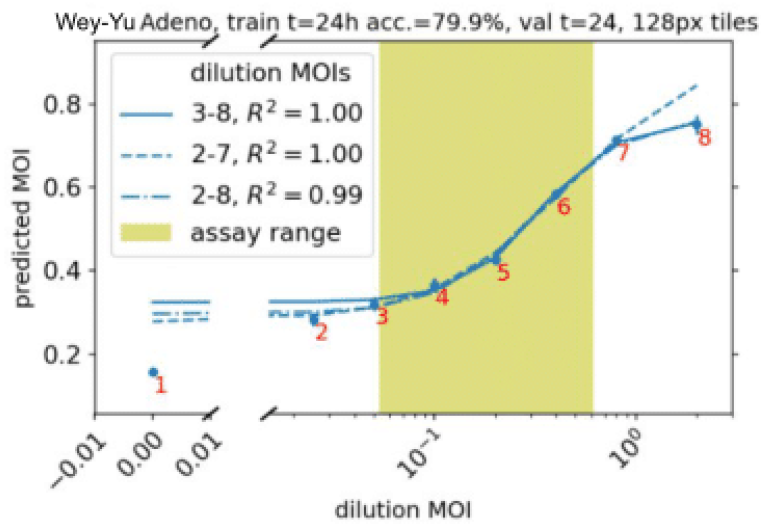
The trained model’s predicted infectivity on the unseen dilution series grouped according to MOI. The displayed error bars are the standard error. The legend displays the dilutions used for the fits and each corresponding goodness of fit score

### 4 Summary

In this report we demonstrated our Al-based model could quantify viral titer on Adenovirus infected HeLa cells in the range 0.053 - 0.61 MOI at a time of 24 hours after initial infection The learning curve and associated confusion matrix demonstrate good training of the Al with the potential for further improvements by increasing the scope of the training data.

## Appendix

### 5 Brightfield Samples

Figure 4 visually demonstrates the quality of the images and the degree of uniformity across the plate. These images indicate that morphological changes in the cells at the different dilutions are not readily apparent by visual inspection at this time point after infection.

**Figure 4:**
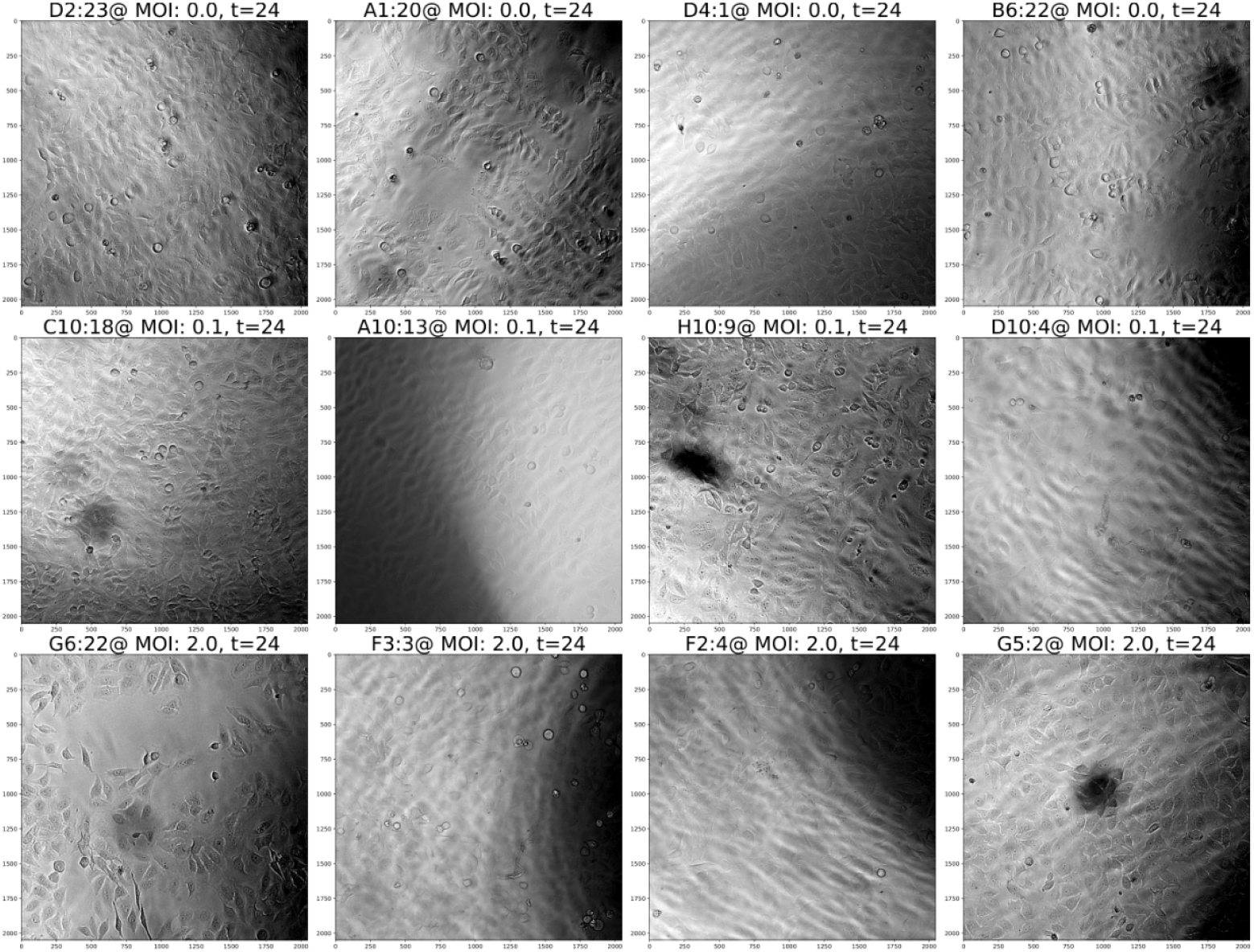
Randomly selected images at several MOIs across the full range. Each image is mean subtracted and scaled to one standard deviation. Above each image is a title containing the well address. MOI and time after infection in hours. These images indicate that morphological changes in the cells at the different dilutions are not readily apparent by visual inspection at this time point after infection.

**Fig. S17. An example training report on adenovirus-infected HeLa cells**. True laboratory identity redacted

